# Evolutionary trajectories of teleost olfactory signaling genes shaped by long-term redundancy after whole-genome duplication

**DOI:** 10.64898/2026.02.17.706259

**Authors:** Tatsuki Nagasawa, Hanami Fujisaki, Takahiro Ogo, Masato Nikaido

**Affiliations:** School of Life Science and Technology, Institute of Science Tokyo, Meguro-ku, Tokyo 152-8550, Japan

## Abstract

Whole-genome duplication (WGD) is a major evolutionary event that drives molecular and species diversification. However, few studies have traced how WGD has shaped the long-term functional evolution of individual genes. Here, we investigated the olfactory marker protein (*omp*) genes duplicated by the teleost-specific WGD (∼300 million years ago) through phylogenetic, syntenic, expression, and promoter analyses. Our results suggest that the duplicated *omp* gene pair has retained redundancy over an extended evolutionary period, leading to both non- and sub-functionalization, thereby generating molecular diversity. Moreover, evolutionary analyses of the olfactory signal transduction cascade revealed prolonged redundancy across its components, likely constrained by gene dosage balance. These findings imply that WGD may have introduced unexpected diversity into the entire olfactory signaling machinery of teleosts through dosage-constrained functional divergence.

**Highlights:**

1. Teleost-specific WGD generated duplicated *omp* genes that persisted for ∼300 million years.
2. Extended redundancy in *ompa*/*ompb* led to both non- and sub-functionalization.
3. Olfactory transduction genes also show long-term redundancy shaped by dosage constraints.
4. WGD likely introduced diversification into teleost olfactory signaling via dosage constraints.

## Introduction

Dr. Susumu Ohno proposed a seminal theory for molecular evolution: i.e. “gene duplication is one of the most critical driving forces for molecular evolution ^1^. Immediately after duplication, gene pairs are in a fully redundant state because they share completely identical sequences, expression patterns, and functions. This redundancy enables one copy to preserve the ancestral function, thereby allowing the other to accumulate mutations under relaxed selective constraints. Consequently, in most duplicated gene pairs, one copy is eventually lost through pseudogenization (non-functionalization), returning to a single-copy state. However, a minority of pairs escape this fate and instead undergo neo-functionalization (acquiring of novel functions) or sub-functionalization (partitioning of ancestral functions). Such divergence has been linked to the acquisition of novel traits ^2^ ^3^ and adaptation to new ecological niches ^4^ ^5^, and has therefore positioned gene duplication as a key mechanism underlying evolutionary innovation. In particular, whole-genome duplication (WGD) is a transformative evolutionary event, creating duplicates of the entire coding gene repertoire and reshaping genome architecture. Independent WGD events have occurred in major lineages such as plants ^67^, yeast ^8,9^, and vertebrates ^10,11,12,13^, and are widely regarded as a source of explosive adaptive radiation ^14^.

Teleost fishes—the largest and most diverse vertebrate group—also experienced an additional WGD approximately 300 million years ago (MYA), known as the teleost-specific or third-round (3R) WGD ^15,16,17,18^. This duplication followed two earlier rounds of genome duplication: the first (1R) in the ancestor of vertebrates and the second (2R) in the ancestor of jawed vertebrates ^19,20^. The teleost-WGD contributed to the evolution of specialized morphologies such as myogenic electric organs ^21^ and the bulbus arteriosus ^22^, facilitating the remarkable ecological expansion of teleosts across diverse aquatic environments. Comparative genomic analyses have revealed that WGD-derived gene pairs follow a two-phase evolutionary trajectory ^23,24^. In the first, rapid phase, 70–80% of duplicates are lost within ∼60 million years (MY) through large-scale chromosomal rearrangements and deletions. The second, prolonged phase involves the gradual accumulation of mutations in the remaining pairs, resulting in slow gene loss and functional divergence. Consequently, prolonged redundancy during the second phase constitutes a critical period for sub-and neo-functionalization. Teleosts thus provide a powerful model for tracing the long-term trajectory of duplicate genes. However, few studies have followed in detail how specific gene pairs avoided early loss and subsequently diversified over hundreds of millions of years.

Olfactory marker protein (OMP) is strongly and specifically expressed in all mature olfactory sensory neurons (OSNs) in mammals, making it a widely used cellular marker ^25,26,27^. Despite its long history as a marker, the molecular function of OMP remained unclear for decades. Recent studies, however, have revealed roles for OMP in calcium homeostasis ^28^, in regulating OSN axonal projection to the olfactory bulb ^29^, and in controlling olfactory adaptation through buffering of the second messenger cAMP ^30^. In teleosts, the OMP gene underwent duplication via the 3R-WGD, giving rise to *ompa* (OMP2) and *ompb* (OMP1). In our previous study ^31^, we demonstrated that *ompa* and *ompb* show clear spatial sub-functionalization in zebrafish. The *ompa* is expressed in OSNs on the apical side of the olfactory lamella, whereas ompb is restricted to basal OSNs, resulting in largely non-overlapping neuronal populations. In addition, the spatial expression of the two genes can be directly compared within the same olfactory epithelial sections. Therefore, teleost *omp* genes serve as an ideal model for long-term functional divergence of duplicates after WGD.

However, previous studies were limited in taxonomic sampling—both in genome surveys and in comparative expression analyses—and lacked data from basal ray-finned fishes (non-teleosts). As a result, the comprehensive evolutionary trajectory of the *omp* gene family remains unresolved. In this study, we expanded the taxonomic breadth of genome mining and spatial expression analyses, and combined these with promoter activity assays to provide a higher-resolution reconstruction of post-WGD evolution of *omp* genes. We show that the duplicated *omp* genes accumulated mutations over extended evolutionary periods and diversified into lineage-specific functionalized states. Furthermore, evolutionary analyses of the entire olfactory signal transduction cascade revealed that all components retained redundancy for hundreds of millions of years, suggesting that long-lasting dosage constraints may have introduced unexpected genetic diversity into the olfactory machinery of teleosts.

## Results

### Non-functionalization of omp genes after WGD

First, we retrieved *omp* gene sequences from whole-genome assemblies of a broad range of representative vertebrates (Table S1). Phylogenetic analyses indicated that teleost *omp* genes were separated into two major clades *ompa* and *ompb*, with moderate bootstrap support for *ompa* (87; Figs. 1A and S1A). To further validate the classification of *omp* genes, we also analyzed the phylogenies of the *capn5* and *myo7a* genes, as *omp* is nested within the second intron of *capn5* ^31^, and *myo7a* is located adjacent to *capn5*. Similar to *omp*, *capn5* and *myo7a* genes were divided into *capn5a*/*capn5b* and *myo7aa*/*myo7ab*, respectively, with strong bootstrap support (100) for the monophyly of both *myo7aa* and *myo7ab* (Figs. 1B–C and S1B–C). We also compared the genomic synteny surrounding *omp* genes (Fig. 1D; details in Fig. S2). In teleosts, as previously reported ^31^, *omp* genes are located in an antisense orientation within the second intron of *capn5*, with the exception of cartilaginous fishes. Genomic synteny around *omp* was well conserved across analyzed species. Non-teleosts generally retained a single-copy synteny set, whereas teleosts maintained two sets on different chromosomes (Figs. 1D and S2). These duplicated synteny sets are characteristic of the teleost WGD and are consistent with our previous study suggesting WGD-mediated duplication of omp genes ^31^. Interestingly, some teleost lineages have lost the *ompb* gene, even though their host *capn5* genes are highly conserved. In addition to phylogenetic and synteny analyses, structural features—such as the conversion of *ompa* into a two-exonic gene by intron insertion—indicate that *ompa* has been retained in all examined species, whereas *ompb* has independently become pseudogenized in several lineages (summarized in Fig. 1E). Supporting this conclusion, fragments of pseudogenized *ompb* were detected in the second intron of *capn5b* in herring and piranha (Fig. S3), suggesting that *ompb* was repeatedly lost through lineage-specific accumulation of mutations, namely non-functionalization. In Cypriniformes (e.g., zebrafish and carp), the full-length *ompb* sequence is preserved, whereas in Characiformes, Siluriformes, and Gymnotiformes, *ompb* has become pseudogenized. Because Cypriniformes and Characiformes diverged approximately 100 million years after the teleost WGD ^32^, this pattern indicates that *ompb* genes survived the initial rapid-loss phase (∼60 MY) and subsequently experienced lineage-specific loss. Taken together, these results demonstrate that although *omp* genes were duplicated in the teleost WGD, lineage-specific non-functionalization of *ompb* generated copy number diversity across species. Despite these changes, cAMP-binding motifs—predicted to mediate the buffering function of OMP ^33,30^—are highly conserved in both *ompa* and *ompb* across all species analyzed (Fig. S4). This conservation suggests that the cAMP-buffering capacity of the duplicated genes has been largely maintained after WGD.

**Figure 1.**
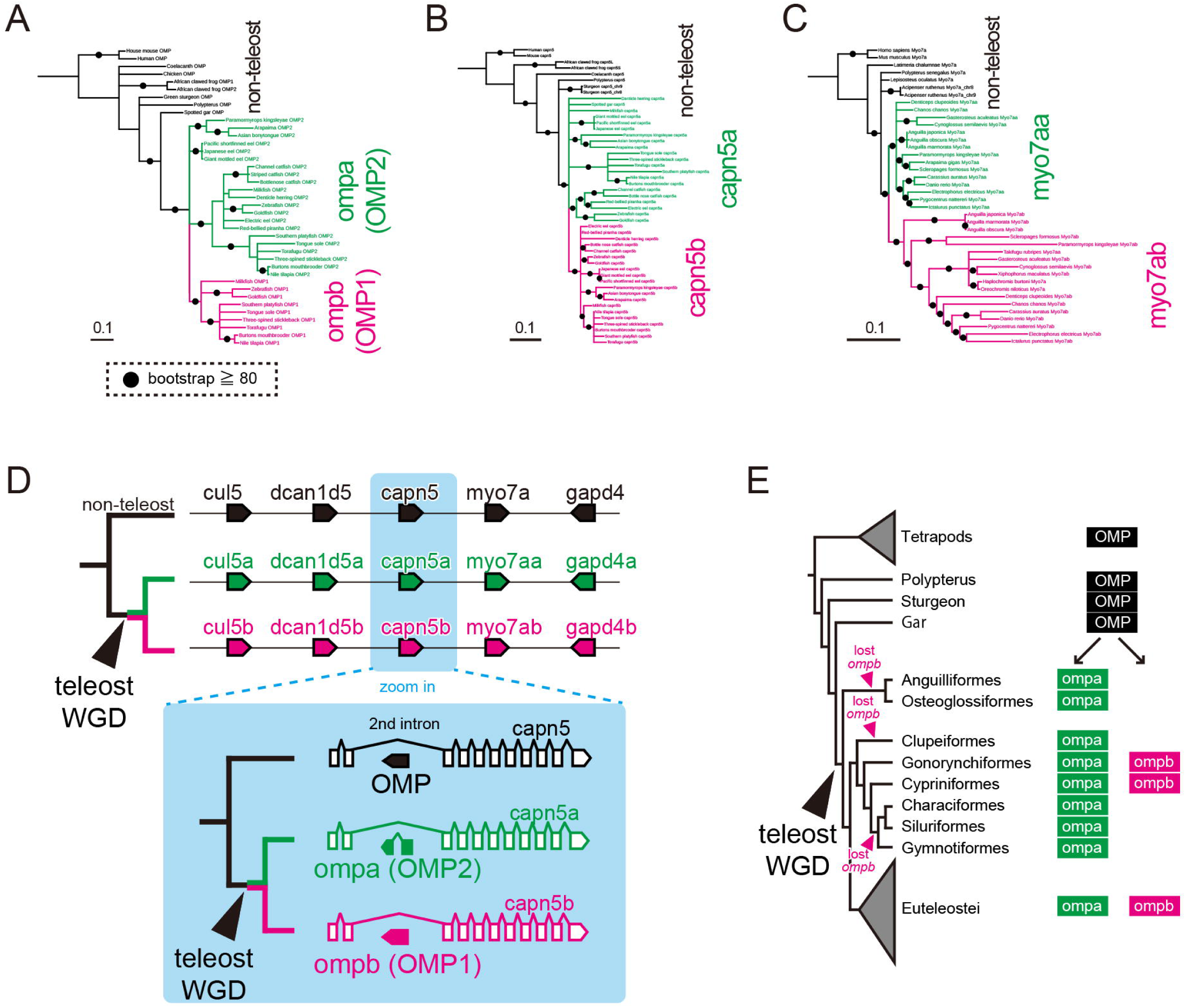
Molecular evolution of the omp gene following teleost whole-genome duplication (WGD). Maximum-likelihood (ML) phylogenetic trees of (A) *omp*, (B) *capn5*, and (C) *myo7a* genes in vertebrates (detailed phylogenetic trees are shown in Fig. S1). Bootstrap values were calculated from 100 replicates; nodes supported by ≥80% bootstrap are marked with black circles, whereas those with <50% support are shown as multifurcations. (D) Genomic synteny and schematic representation of the *omp* gene and its neighboring genes (detailed synteny information is provided in Fig. S2). The *omp* gene is a nested gene located in the reverse orientation within the second intron of *capn5*. The genomic synteny of *omp* and its neighboring genes (*cul5*, *dcan1d5*, *capn5*, *myo7a*, and *gapd4*) is highly conserved across vertebrates. In teleosts, two sets of these genes are located on different chromosomes, consistent with their origin as WGD-derived duplicates. Notably, in all teleost species examined, the *ompa* gene contains a single intron that divides the coding region. (E) Schematic representation of the molecular evolution of *omp* genes. In non-teleost ray-finned fishes (e.g., *Polypterus*, sturgeon, and gar), *omp* is coded as a single-copy gene (black). Following the teleost WGD, *omp* duplicated into *ompa* (green) and *ompb* (magenta). While *ompa* was retained in all teleost lineages analyzed, *ompb* was secondarily lost in multiple lineages (indicated by magenta arrowheads).

### Effects of WGD and pseudogenization on spatial expression patterns

To infer how WGD-driven duplication and subsequent pseudogenization affected *omp* gene expression, we performed fluorescent *in situ* hybridization to visualize spatial expression patterns in six representative fish species (Fig. 2). These species were grouped into three categories: (1) non-WGD species (Senegal bichir *Polypterus senegalus* and spotted gar *Lepisosteus oculatus*), (2) teleosts retaining both *omp* genes after WGD (zebrafish *Danio rerio* and African cichlid *Haplochromis sauvagei*), and (3) teleosts that lost *ompb* after WGD (Japanese eel *Anguilla japonica* and red-bellied piranha *Pygocentrus nattereri*). For each category, two phylogenetically distant species were analyzed. We first examined expression in retinal horizontal cells (hc). Consistent with previous studies in zebrafish ^31^, gene expression was detected in the horizontal cells of all analyzed species. In non-teleosts, the single-copy omp gene was expressed, whereas in WGD-experienced teleosts, only the *ompa* gene was expressed in horizontal cells. These results suggest that expression in horizontal cells was maintained in *ompa* via sub-functionalization after WGD. In contrast, *capn5*, the host gene of *omp*, was expressed not in horizontal cells but predominantly in the inner nuclear layer (Fig. S5).

**Figure 2.**
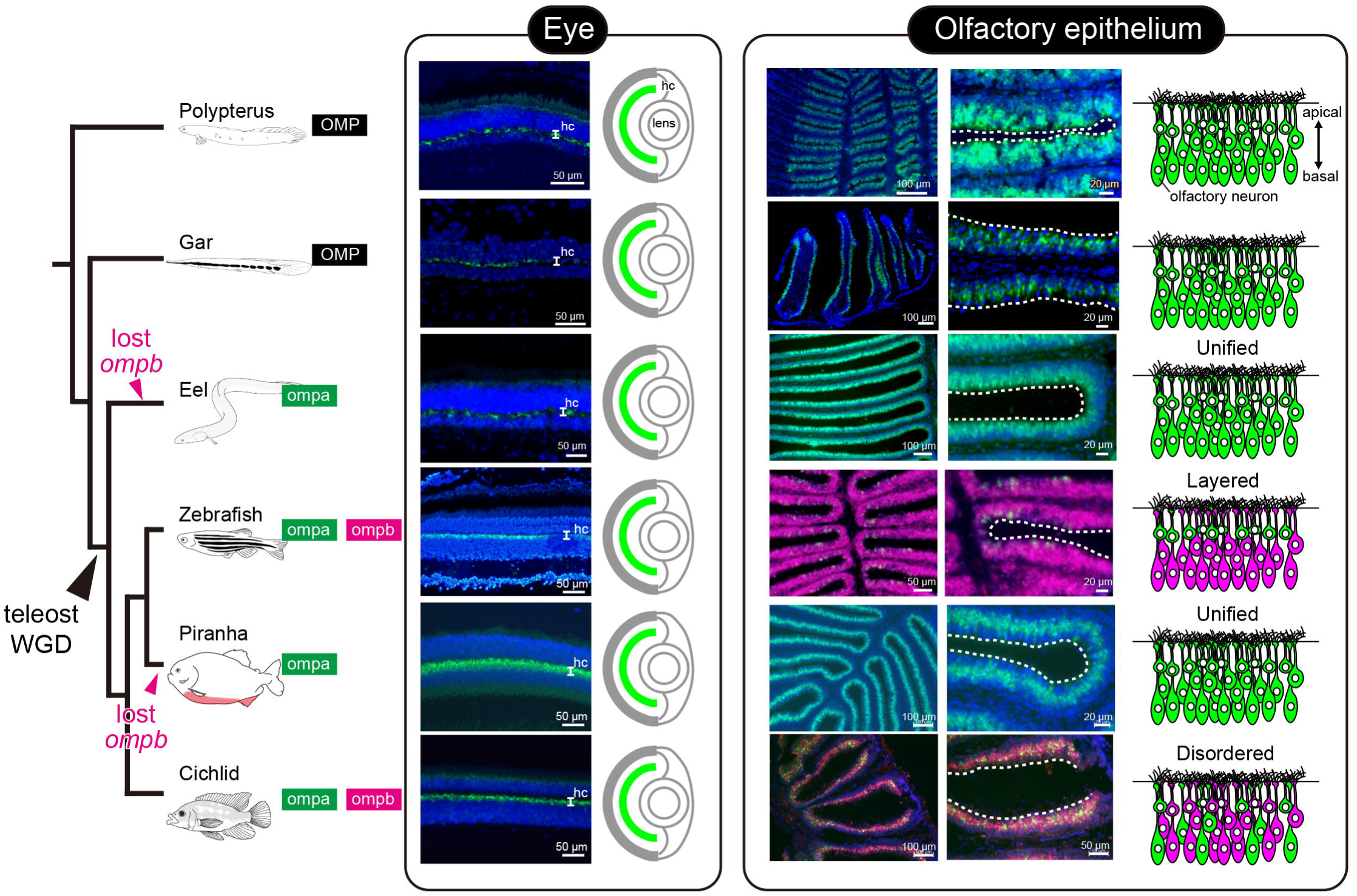
Evolution of the spatial expression patterns of *omp* genes at the cellular level. The phylogenetic tree on the left illustrates the evolutionary relationships of the six species analyzed in this study, along with the *omp* genes they possess and an overview of their evolutionary history. The central panels show expression in the eye, particularly in horizontal cells (hc) of the retina, along with a schematic representation. The right panels depict expression in the olfactory epithelium and its corresponding schematic.

Next, we examined expression in the olfactory epithelium (Fig. 2). In non-teleosts (*Polypterus* and spotted gar), the single-copy *omp* gene was broadly expressed across the olfactory epithelium, similar to mammals. In teleosts retaining both *ompa* and *ompb* (zebrafish and cichlid), both genes were expressed in the olfactory epithelium. In zebrafish, as previously reported, *ompa* and *ompb* exhibited largely mutually exclusive expression, with *ompa* localized to the apical side of the olfactory lamellae and *ompb* to the basal side; the number of *ompb*-expressing cells was predominant. In cichlid, *ompa* and *ompb* were also expressed mutually exclusively, with *ompb*-expressing cells predominating, although a clear layered expression pattern was not observed as in zebrafish. In teleosts that lost *ompb* (eel and piranha), *ompa* was expressed across the entire population of olfactory sensory neurons. These results suggest that each fish species maximally utilizes its available set of *omp* genes, expressing them throughout the olfactory sensory neuron population in a species-specific manner.

### Evaluation of promoter activity across omp genes

To investigate the mechanisms underlying the diversification of spatial expression patterns among *omp* genes, we generated zebrafish reporter lines driven by promoter sequences from various species (Fig. 3). All reporter lines exhibited clear fluorescent signals in the olfactory organ at the larval stage (Fig. 3A–F, A’–F’). In adult olfactory epithelium (OE), the zebrafish *ompa* promoter drove Venus expression in the apical layer (Fig. 3G, G’), whereas the *ompb* promoter drove expression in the basal layer of the olfactory lamellae (Fig. 3H, H’). These expression layers closely mirrored the endogenous mRNA patterns, indicating that each promoter accurately captures the transcriptional properties of its corresponding *omp* gene. In contrast, promoter sequences from species that did not experience the teleost WGD—the spotted gar (Fig. 3I and I’) and mouse (Fig. 3J and J’)—drove Venus expression across both layers, even in a different species (zebrafish OE). Likewise, promoters derived from cichlid *omp* genes produced disordered expression patterns (Fig. 3K, K’, L, L’) that mirrored the original cichlid OE patterns, rather than the layered organization characteristic of zebrafish *ompa* and *ompb*. Similarly, in the adult retina, Venus expression driven by each promoter consistently reflected the endogenous expression layers of the corresponding genes (Fig. 3M–R). Taken together, these results indicate that *omp* promoters from diverse species can reproduce their species-specific expression patterns even in a different species (zebrafish) cellular environment. This pattern strongly supports the view that the diversification of *omp* expression among teleosts has been shaped primarily by mutations accumulated within cis-regulatory elements, rather than by shifts in trans-regulatory factors, chromatin accessibility, or the spatial arrangement of olfactory sensory neuron subtypes. A summary of the molecular evolutionary patterns revealed in this study is provided in Fig. 4.

**Figure 3.**
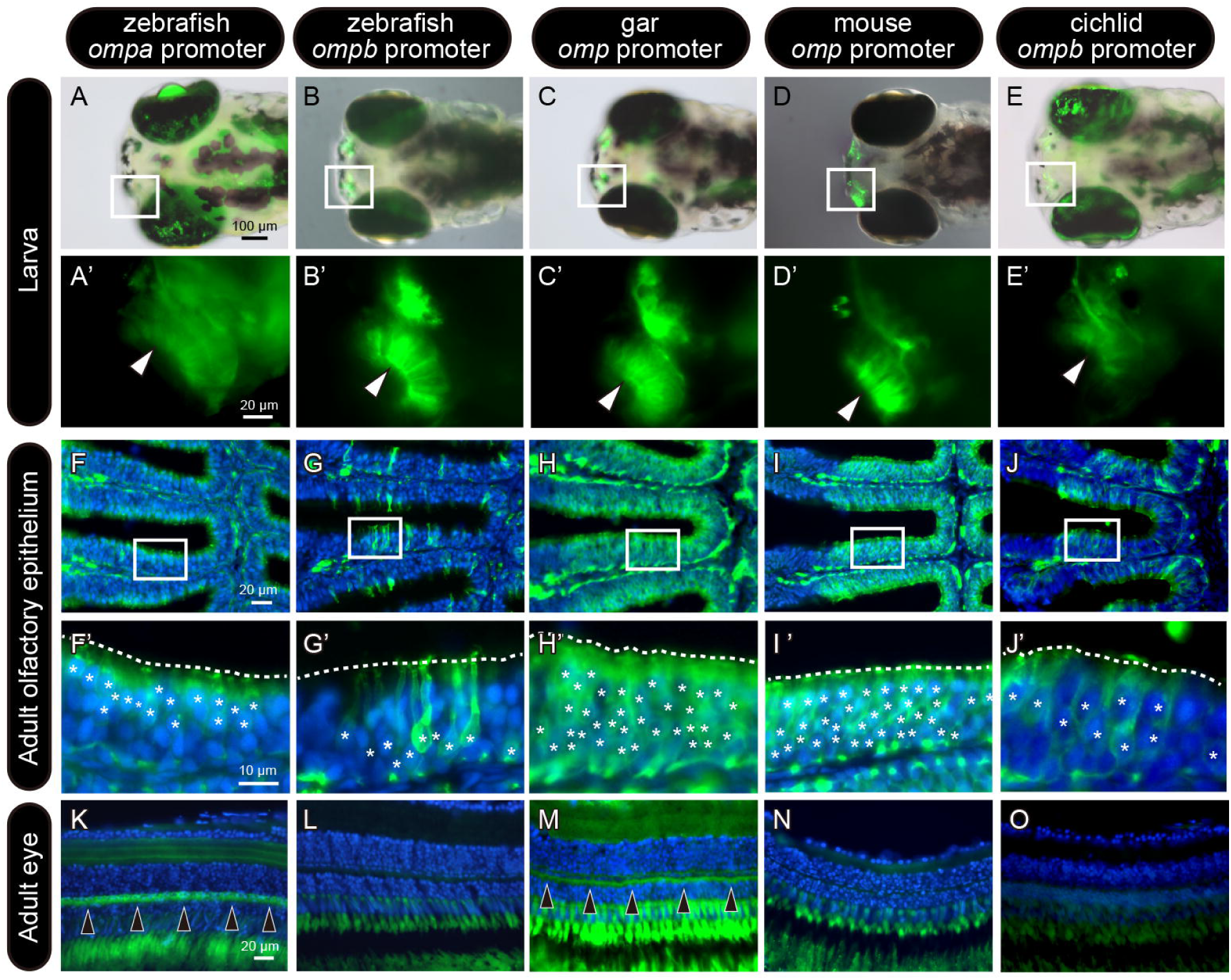
Reporter assay of *omp* promoter activity using transgenic zebrafish lines. (A–E) Reporter fluorescence in 5 dpf (days post-fertilization) larvae of transgenic zebrafish. Each panel shows an overlay of bright-field and fluorescent images. (A′–E′) High-magnification images of the olfactory organ in the corresponding transgenic zebrafish larvae shown in panels A–E. (F–J) Sections of the olfactory epithelium in adult transgenic zebrafish lines. The boxed regions are shown at higher magnification in panels F′–J′. (K–O) Sections of the eye in adult transgenic zebrafish lines. The promoters of zebrafish *ompa* and gar *omp* drove reporter expression in horizontal cells of the retina (arrowheads).

**Figure 4.**
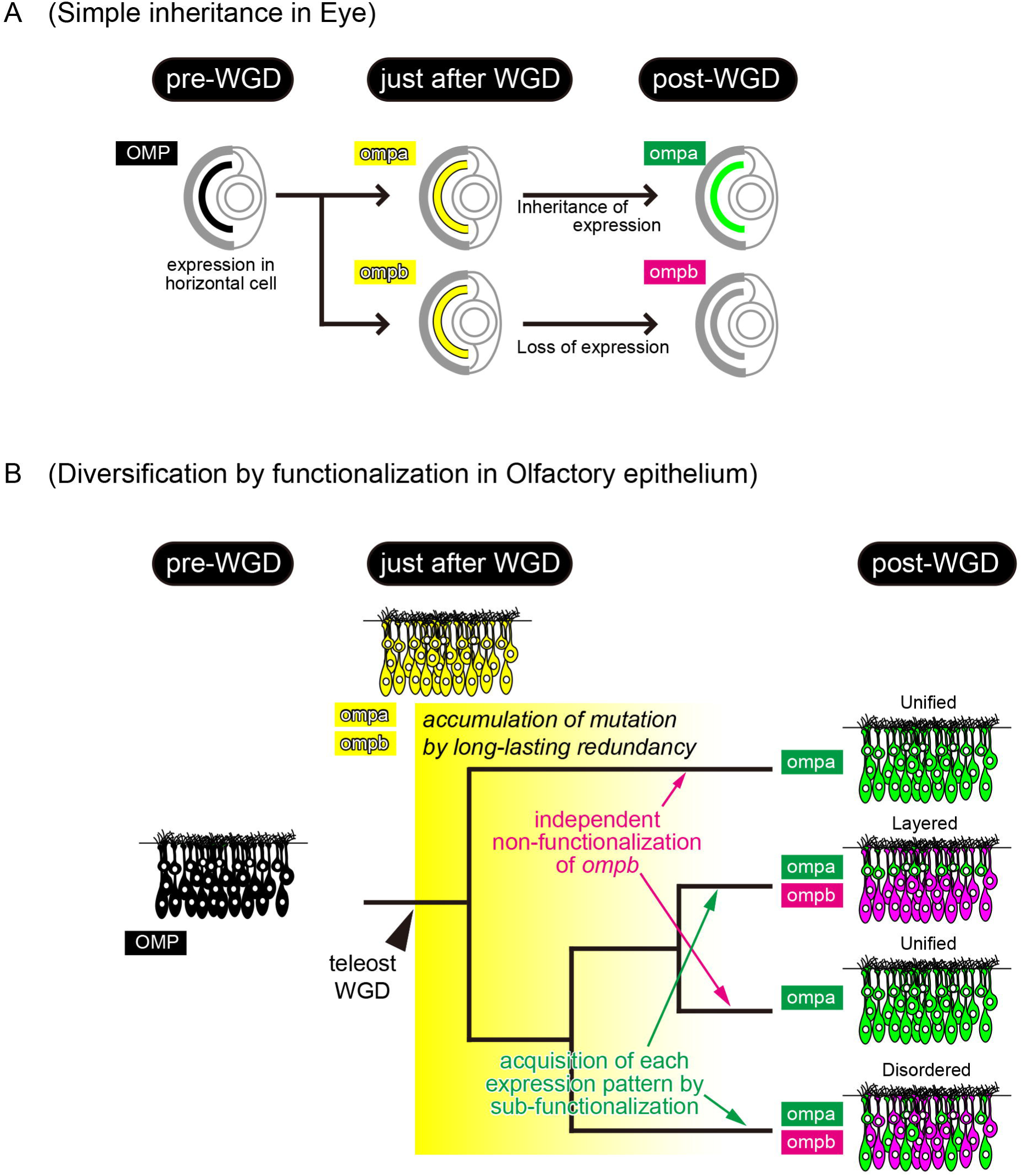
Schematic representation of the impact of teleost WGD on the spatial expression patterns of *omp* genes. (A) Evolutionary changes in *omp* expression in horizontal cells of the eye — functionalization by simple inheritance of the original expression pattern in one of the duplicated genes. Before the teleost WGD, *omp* was expressed in retinal horizontal cells (before WGD). Immediately after the WGD, both *ompa* and *ompb* were likely expressed in horizontal cells (just after WGD). Over the long course of evolution, *ompa* retained expression in horizontal cells, whereas *ompb* lost this expression (after WGD). (B) Evolutionary changes in *omp* expression in olfactory sensory neurons — diversification of expression patterns through accumulation of mutations under long-lasting redundancy. Prior to the teleost WGD, *omp* was expressed throughout mature olfactory sensory neurons (before WGD). Immediately after WGD, both *ompa* and *ompb* were likely expressed throughout mature olfactory neurons (just after WGD). Over evolutionary time, the redundancy maintained by the two pairs of genes (*ompa* and *ompb*) allowed accumulation of cis-regulatory mutations, leading to diversification of expression patterns during species divergence (after WGD).

### Dosage constraints and long-lasting redundancy in olfactory signaling cascade

Inoue et al. estimated that approximately 72–82% of WGD-derived duplicate pairs experienced loss of one paralog within the first ∼60 MY ^24^. To clarify how *omp* genes escaped this early phase of extensive gene loss, we analyzed the molecular evolutionary patterns of genes involved in the olfactory signal transduction cascade (Fig. 5). Unlike small-scale duplications, WGD duplicates entire interacting gene networks. As a result, overall dosage balance is preserved even as copy numbers increase. In a dosage-sensitive network, the loss of even a subset of components disrupts functional stoichiometry, thereby constraining the loss of duplicate genes (Fig. 5A; the dosage-constraints hypothesis, also referred to as dosage-balance or dosage-compensation) ^34,35,36^. The vertebrate olfactory signal transduction cascade—particularly well characterized in mouse—includes the genes *gnal, adcy3, omp, cnga2, ano2,* and *slc24a4* (Fig.5B)^37,38,39,40,41^. Using whole-genome searches, phylogenetic analyses, and genomic synteny comparisons, we identified the orthologs and WGD-derived paralogs of these six genes across teleost lineages (Fig. S6). Strikingly, in many teleost clades, both WGD-derived pairs of all six genes have been retained (Fig. 5C). These findings indicate that, in contrast to the genome-wide trend reported by a previous study ^24^, in which roughly 70–80% of gene duplicates were lost during the first ∼60 MY after WGD, olfactory signaling genes exhibit exceptional long-term retention. In contrast, the olfactory transduction cascade exhibits a distinct tendency toward long-term retention of WGD-derived paralogs.

**Figure 5.**
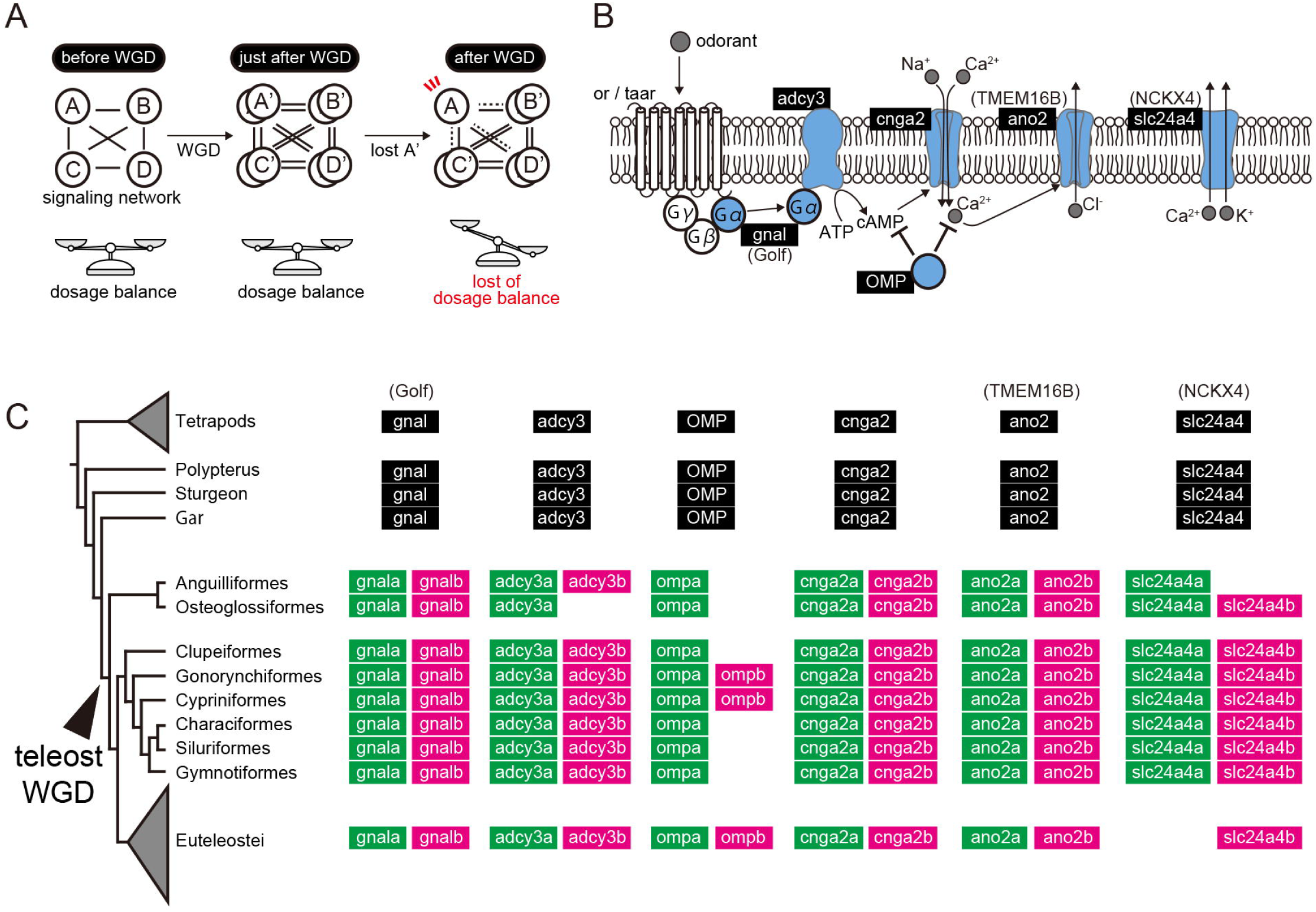
Dosage constraints hypothesis in the olfactory signaling cascade following teleost WGD. (A) Schematic illustration of the dosage constraints hypothesis. At the cellular level, genes constituting a signaling network or metabolic pathway maintain appropriate relative levels of translated proteins (before WGD). WGD doubles all genes, thereby proportionally increasing the amount of each protein while maintaining their relative balance (just after WGD). Subsequently, if one of the duplicated genes in the network becomes pseudogenized, the stoichiometric balance of protein components is disrupted, leading to network collapse (after WGD). Because individuals with a disrupted network are eliminated by purifying selection, the WGD-derived gene pairs tend to be retained over evolutionary time — a process explained by the dosage constraints hypothesis. (B) Schematic diagram of a representative vertebrate olfactory signaling cascade. (C) Molecular evolution of olfactory signaling cascade genes before and after teleost WGD, showing the presence or absence of each gene across lineages, based on phylogenetic and genomic synteny analyses (see also Figure S5).

## Discussion

Gene duplication is a major driver of molecular evolution, and, in particular, whole-genome duplication (WGD), which duplicates the entire gene repertoire, represents a prominent example ^1^. While WGD can facilitate the acquisition of novel traits and diversification of morphology and function ^21,22,14^, only a small subset of duplicated genes actually contributes to these innovations. The majority of duplicated gene pairs are thought to rapidly revert to a single-copy state due to functional redundancy^23,24^. Nevertheless, many gene pairs retain long-term redundancy and undergo lineage-specific functional divergence, including vertebrate Gnrh1/Gnrh3 ^42^, teleost cholecystokinin a/b ^43,44^, yeast ribosomal protein genes ^45^, and plant SEP1/SEP2 and SHP1/SHP2 ^46,47^. These cases illustrate that WGD-derived genes do not necessarily undergo rapid divergence or loss but can follow diverse evolutionary trajectories through prolonged redundancy.

Teleosts, which experienced WGD approximately 300 million years ago (MYA), provide an ideal model to investigate the long-term molecular evolution of duplicated genes ^23,24^. However, studies that integrate multiple aspects—such as gene sequence, expression patterns, and transcriptional regulation—to trace the long-term evolutionary trajectory following WGD remain limited. Here, we focused on the teleost WGD-derived *omp* (olfactory marker protein) genes to explore diverse trajectories of gene functional divergence over ∼300 million years (MY). By combining genomic analyses, comparative expression studies, and promoter activity assays, we showed that duplicated *omp* gene pairs have undergone lineage-specific accumulation of mutations, resulting in either sub-functionalization or non-functionalization. As illustrated in Fig. 4, *ompa* and *ompb* likely shared ancestral expression patterns immediately after WGD. Subsequently, expression in the visual system was largely and directly inherited by *ompa*, whereas in the olfactory system, lineage-specific mutations led to diverse expression patterns, resulting in sub-functionalization or non-functionalization. These results suggest that functionalization in the olfactory system was not a simple inheritance of ancestral function but rather the outcome of a complex and dynamic molecular evolutionary process.

### Functional role and evolutionary origin of omp

Our study revealed that WGD-derived *ompa*/*ompb* pairs have been largely retained in many teleosts, and the two genes showed clear transcriptional differentiation in zebrafish and cichlid (Fig. 2). Moreover, *ompa* expression in horizontal cells of the retina is conserved in teleosts, and similar expression is observed in basal ray-finned fishes (*Polypterus* and gar) that retain a single-copy *omp* gene. These patterns indicate sub-functionalization, although physiological divergence between ompa and ompb remains unclear. Previous studies have shown that the cAMP-binding motif of the omp protein is highly conserved from mammals to fish ^33,30^. In our analysis distinguishing between *ompa* and *ompb*, this motif was highly conserved in both, preventing functional differentiation from being inferred based on sequence alone (Fig. S4).

The *omp* is found in both bony fishes and tetrapods, and its origin has been suggested to date back to at least the common ancestor of bony fishes ^31^. In this study, we confirmed that *omp* is absent from the introns of *capn5* in cartilaginous fishes, strongly supporting that *omp* was newly acquired in the bony fish lineage (Fig. S2). Furthermore, *omp* expression in horizontal cells suggests that this trait predates the common ancestor of ray-finned fishes (Fig. 2). However, *omp* expression in the retina is not detected in mouse ^31^, and data from basal sarcopterygians such as lungfish are limited, leaving the precise timing of acquisition of horizontal cell expression unresolved. Given that jawless vertebrates are known to possess horizontal cells ^48,49,50^, it is likely that *omp* function in horizontal cells evolved independently of the acquisition of horizontal cells during vertebrate evolution.

The apparent restriction of *omp* expression in horizontal cells to ray-finned fishes may reflect visual adaptations to aquatic environments. In aquatic environments, turbidity reduces light transmission and shifts spectral composition dynamically, making rapid and flexible light adaptation essential ^51,52^. Horizontal cells in teleosts are strongly modulated by dopamine via the D1 receptor–cAMP–PKA pathway, which regulates gap-junction coupling and light responsiveness ^53,54^. Because *omp* buffers intracellular cAMP levels to regulate sensory adaptation in olfactory neurons ^30^, we hypothesize that *ompa* in horizontal cells may similarly stabilize cAMP dynamics to fine-tune light-adaptation kinetics. Such modulation could help maintain visual performance under dynamic underwater light environments. Future knockout studies will be needed to determine its precise physiological function.

### Regulation of omp gene expression

The *omp* gene is a nested gene located within an intron of the *capn5* host gene (Fig. 1D). The regulatory relationship between host and nested genes—whether they are co-expressed, mutually exclusive, or independent—has been debated for decades ^55,56,57^. Previous studies in mouse showed that in olfactory sensory neurons (OSNs) where the nested *omp* gene is expressed, the host gene *capn5* is not expressed ^58^. In the present study, analysis of capn5 expression in the retina of gar and zebrafish revealed expression in the inner nuclear layer, whereas no signal was detected in horizontal cells expressing *omp* (Fig. S5). This separation of expression suggests that the *omp* gene is regulated independently of its host gene. In mammals, short promoter regions of approximately 0.3 kb are sufficient to drive mature OSN-specific expression ^59,60,61^, and in the present study, upstream sequences of ∼1 kb mimicked endogenous expression patterns. These observations indicate that *omp* transcription is driven by a short upstream sequence across vertebrates and is regulated independently of the host gene.

### Dosage constraints promote maintenance and functional divergence of teleost olfactory genes

An important question in molecular evolution concerns why some WGD-derived gene pairs, such as *ompa*/*ompb*, have been retained over long evolutionary periods. In this study, we hypothesized that dosage constraints within olfactory sensory neurons have promoted the maintenance of *ompa*/*ompb* gene pairs. According to the dosage-constraints hypothesis ^34,35,36^, maintaining the quantitative balance of component genes within molecular complexes functioning in the cell is advantageous under natural selection. Consistent with this hypothesis, Sato et al. reported that gene pairs retained long-term after the teleost-WGD tend to have numerous interacting partners^23^. In this study, we analyzed multiple genes comprising the olfactory transduction cascade (*gnal*, *adcy3*, *cnga2*, *ano2*/*tmem16b*, *slc24a4b*) and found that all duplicated gene pairs are retained in teleosts (Fig. 5). This retention is remarkable considering that 72–82% of gene pairs were lost during the first ∼60 MY following the teleost-WGD ^24^ and that the retention rate of WGD-derived gene pairs in zebrafish is only ∼26% ^62^. These observations imply that strong dosage constraints act on the olfactory pathway, maintaining functional redundancy. Moreover, prolonged maintenance of redundancy facilitates neo- or sub-functionalization ^63^. The partial co-expression of *ompa* and *ompb* in a subset of olfactory sensory neurons (Fig. 2) ^31^ may reflect this transitional state. Taken together, long-term redundancy after WGD, followed by subsequent functional differentiation, likely promoted diversification of the olfactory transduction cascade and provided an evolutionary foundation for teleosts to adapt to diverse olfactory environments. A more detailed understanding of the divergence and evolutionary trajectories of olfactory signaling molecules in fishes remains an important future goal.

## Methods

### Animals and ethical approval

Adult Spotted gar (*Lepisosteus oculatus*), Gray bichir (*Polypterus senegalus*), Japanese eel (*Anguilla japonica*), Red-bellied piranha (*Pygocentrus nattereri*), and African cichlid (*Haplochromis sauvagei*) were obtained from commercial suppliers. Zebrafish (*Danio rerio*) of the inbred RIKEN Wild type strain, provided by the National BioResource Project (NBRP, Japan), were used in this study. All fish were maintained in breeding tanks at 25–29 °C under a 12 h light / 12 h dark photoperiod and fed two to three times daily. For tissue sampling, fish were anesthetized with MS-222 and euthanized by rapid decapitation, after which the olfactory rosettes and eyes were extracted. Genomic DNA for amplification of the mouse promoter sequence was kindly provided by Prof. Junji Hirota (Institute of Science Tokyo). Accordingly, no live mouse experiments were conducted in this study. All recombinant DNA experiments were approved by the institutional biosafety committee of the Institute of Science Tokyo (I2019035 and I2024021).

### Isolation and collection of gene sequences

The identification and isolation of gene sequences were performed following previously published methods ^64^ in accordance with slight modifications. Gene sequences were either retrieved from the NCBI database or newly predicted from whole-genome assemblies deposited in NCBI. To identify the target loci, TBLASTN searches were performed against the genomic sequences of each species using amino acid sequences of orthologous genes from closely related species as queries. Subsequently, exon sequences were extracted from the surrounding genomic regions using Genewise2 ^65^, with the same orthologous amino acid sequences as references. All accession numbers and the sequences used in this study are summarized in Table S1.

### Molecular phylogenetic and genomic synteny analysis

Maximum-likelihood phylogenetic analyses were performed using RAxML-NG ^66^. Amino acid sequences of each gene were aligned using MAFFT ^67^, whereas *omp* and *capn5* nucleotide sequences were aligned with MAFFT (nucleotide mode) followed by codon-based alignment. The best-fit substitution models were determined using ModelTest-NG ^68^, and phylogenetic trees were inferred with RAxML-NG with 100 bootstrap replicates. Phylogenetic trees were visualized using iTOL ver. 6 ^69^(https://itol.embl.de/). Genomic synteny comparisons were performed using Genomicus ^70^(https://www.genomicus.bio.ens.psl.eu/genomicus-110.01/cgi-bin/search.pl) and NCBI Genome Data Viewer ^71^(https://www.ncbi.nlm.nih.gov/gdv/). Homologous relationships among genes were carefully validated through TBLASTN searches against the available genome datasets of each species.

### Construction of reporter constructs and establishment of transgenic zebrafish lines

For the establishment of each transgenic zebrafish line, we used the plasmids pT2AL200R150G and pCS-zTP, which were kindly provided by Prof. Koichi Kawakami (National Institute of Genetics) (https://ztrap.nig.ac.jp/trans.html), following the procedures described previously ^72^ with minor modifications. The GFP sequence of pT2AL200R150G was replaced with the Venus fluorescent protein sequence (hereafter referred to as the Venus vector), which was used as the backbone for all reporter constructs. Genomic DNA was used as a template to amplify the upstream region of the omp gene by PCR. The primers used for amplification are listed in Table S3. Because PCR amplification of the upstream region of zebrafish *ompa* was difficult due to repetitive sequences, a DNA fragment excluding the repetitive region was synthesized commercially (Integrated DNA Technologies; the sequence is shown in Figure S6). The amplified fragments were cloned into the Venus vector, and plasmid DNA was obtained after transformation into *Escherichia coli*. The resulting plasmid and transposase mRNA, synthesized from pCS-zTP using SP6 RNA polymerase (mMESSAGE mMACHINE SP6 Transcription Kit; Invitrogen), were co-injected into zebrafish fertilized eggs. Individuals showing fluorescence signals in olfactory sensory neurons at 5 days post fertilization (dpf) were raised as founder (F0) fish. The matured F0 fish were crossed with wild-type individuals, and the offspring showing similar fluorescence signals were selected and maintained as F1. The transgenic lines showing stable fluorescence expression over F2 and subsequent generations were established and used for analyses.

### Preparation of frozen sections and in situ hybridization

Frozen sections were prepared as follows. Tissues were dissected in chilled 4% paraformaldehyde (PFA) and subsequently fixed overnight at 4D. For the eyes, the cornea was carefully punctured with forceps to facilitate the penetration of PFA. After fixation, the samples were thoroughly rinsed in PBS, cryoprotected overnight in 30% sucrose solution, and embedded in OCT compound (SAKURA Finetek Japan). Embedded tissues were sectioned at a thickness of 10 µm using a cryostat, mounted onto glass slides, and either observed under fluorescence microscopy after embedding with VECTASHIELD mounting medium with DAPI (Vector Laboratories) or stored at −80 °C until use for *in situ* hybridization. In order to visualize mRNA localization, *in situ* hybridization was performed according to previous study ^31,73^ with slight modification. Specific primers (listed in Table S3) were designed within the coding region of the *omp* genes, and PCR amplification was performed using olfactory epithelium cDNA as a template. Amplified fragments were cloned into the pGEM-T vector (Promega). The plasmids were linearized by restriction enzyme digestion, and DIG-labeled RNA probes were synthesized using the DIG RNA Labeling Kit (Roche). For dual in situ hybridization, FITC-labeled RNA probes were synthesized in the same manner using the FITC RNA Labeling Mix (Roche). For single-color detection, prior to hybridization, the sections were treated with proteinase K for protein digestion, followed by post-fixation, inactivation of endogenous peroxidase (POD) and alkaline phosphatase (AP), and acetylation of amino groups.

Hybridization was carried out overnight at 55–70 °C. After stringent washing to remove unbound probes, sections were incubated with 1% Blocking Reagent (Roche), followed by anti-digoxigenin-POD Fab fragments (Roche) at 4 °C overnight. Signals were amplified using the TSA Biotin System (Akoya Biosciences), and after blocking endogenous biotin and streptavidin, visualization was performed with Streptavidin-Alexa Fluor 488 conjugate (Invitrogen). Sections were mounted in VECTASHIELD with DAPI for nuclear counterstaining and imaging. For dual-color detection, the procedures were generally identical to those for single-color detection, except that signal amplification was performed using the TSA Plus DIG System (Akoya Biosciences), followed by POD inactivation with HDOD treatment. Sections were then reacted with anti-fluorescein-POD Fab fragments (Roche) and anti-digoxigenin/digoxin antibody DyLight 594 (Vector Laboratories), and further amplified using the TSA Biotin System. Visualization was achieved with Streptavidin-Alexa Fluor 488 (Invitrogen), and the sections were mounted and observed as described above.

## Lead contact

Masato Nikaido

## Acknowledgments

We thank Prof. Koichi Kawakami (National Institute of Genetics, Japan) for kindly providing the *Tol2* plasmid. This study was supported in part by GrantDinDAid for Young Scientists (JSPS KAKENHI Grant Number 23K14249), the Sasakawa Scientific Research Grant from the Japan Science Society to T.N., and Grant in Aid for Scientific Research B (JSPS KAKENHI Grant 24K02074) to M.N.

## Author contributions

All authors wrote, revised and confirmed manuscript. Conceptualization, T.N. and M.N. Expression analysis, T.N. and H.F. Syntenic and Phylogenetic analysis, T.N. and M.N. Establishment of transgenic line, T.N. Supervision, M.N. Funding acquisition, T.N. and M.N.

---

**Figure S1. Detailed maximum-likelihood phylogenetic trees shown in Figure 1.**

Detailed views of the maximum-likelihood phylogenetic trees for (A) *omp*, (B) *capn5*, and (C) *myo7a* shown in Figure 1. Bootstrap values are indicated on each node (values below 50% are shown as multifurcations).

**Figure S2. Genomic synteny of omp genes.**

Schematic representation of the genomic synteny surrounding the *omp* (*capn5*) genes in vertebrates. The orientation of pentagons indicates the transcriptional direction of each gene, and orthologous genes across species are connected with dashed lines. The presence or absence of *omp* within the second intron of *capn5* is highlighted in the dotted box (loss of *omp* is shown as an open pentagon with dashed outlines).

**Figure S3. Fragments of pseudogenized ompb gene.**

(A) Herring and (B) piranha show traces of pseudogenized *ompb* fragments, revealed by translating the nucleotide sequences located within the second intron of *capn5b* and comparing them with *ompb* from the closely related zebrafish. The amino acid sequence of zebrafish *ompb* is shown in the upper panel (boxed), whereas the nucleotide and translated amino acid sequences of herring and piranha are shown below. Identical amino acids are highlighted in magenta and connected by lines.

**Figure S4. Multiple alignment of omp genes and the conserved cAMP-binding motif.**

Multiple alignment of amino acid sequences of all *omp* genes analyzed in this study. Gaps are indicated by dashes, and identical residues are shown in blue. The putative cAMP-binding motif suggested by previous studies ^33,30^ is highlighted with a blue background.

**Figure S5. Expression pattern of the capn5 gene in the retina**

Identification of retinal layers was based on a previous study (Yazulla and Studholme, *J. Neurocytol.* 2002). ONL: outer nuclear layer; OPL: outer plexiform layer; INL: inner nuclear layer; IPL: inner plexiform layer. All *capn5* genes were expressed not in the horizontal cells (hc), where the nested gene *omp* is expressed, but rather in the INL.

**Figure S6. ML phylogenetic trees and genomic synteny of olfactory signaling cascade genes.**

(A and B) gnal, (C and D) adcy3, (E and F) cnga2, (G and H) ano2, and (I and J) slc24a4. Bootstrap values are indicated for all nodes in each phylogenetic tree. Genomic synteny diagrams were illustrated using the same manner as described in Figure S2.

**Figure S7. Modified upstream sequence of zebrafish ompa inserted into the Venus vector.**

The modified upstream sequence of the zebrafish *ompa* gene inserted into the Venus vector is shown in FASTA format.

