## Supplementary material for "Evolutionary trajectories of teleost olfactory signaling genes shaped by long-term redundancy after whole-genome duplication": Suppl_Figures

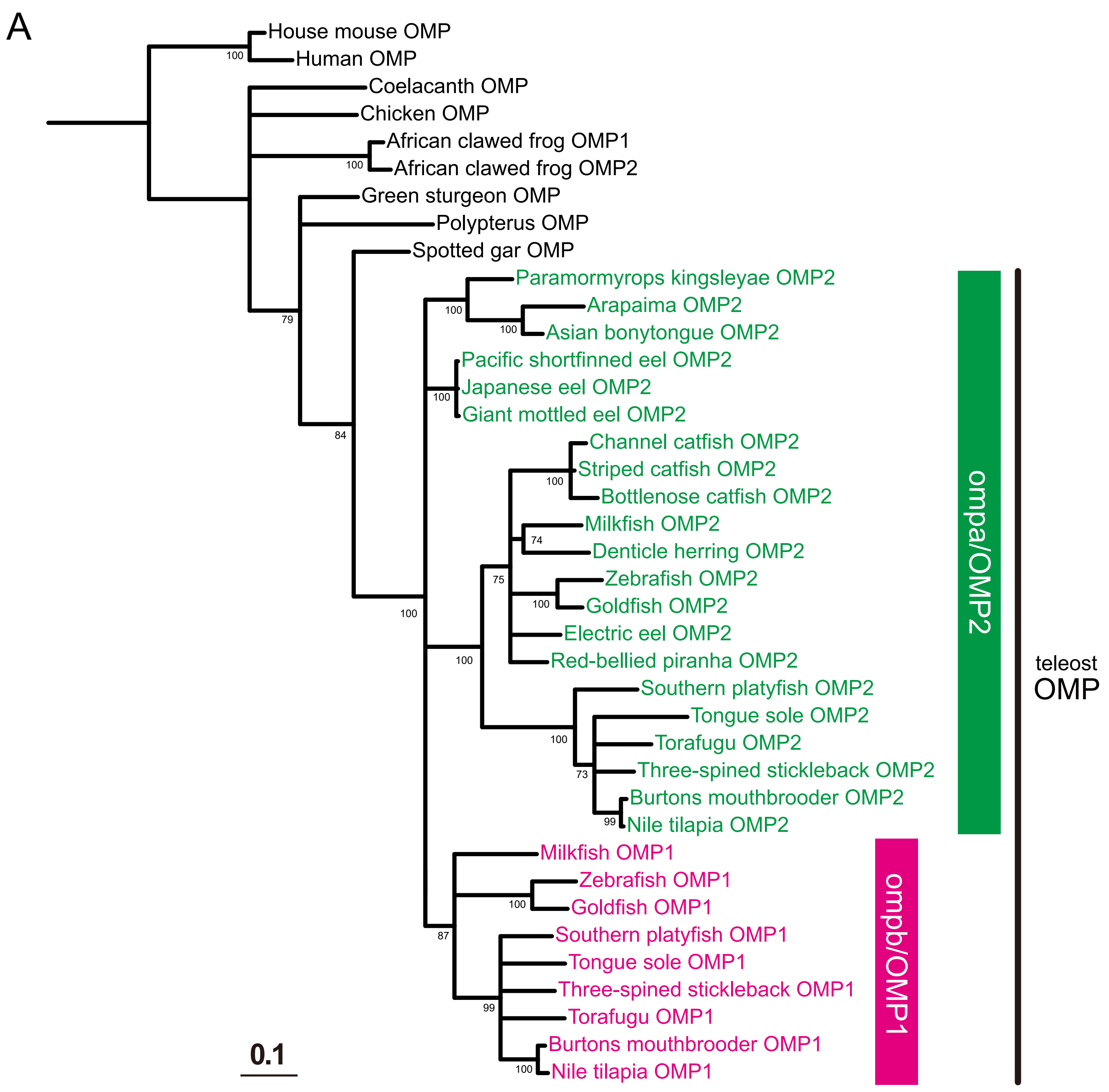
A
House mouse OMP
Human OMP
Coelacanth OMP
Chicken OMP
African clawed frog OMP1
African clawed frog OMP2
Green sturgeon OMP
Polypterus OMP
Spotted gar OMP
Paramormyrops kingsleyae OMP2
Arapaima OMP2
Asian bonytongue OMP2
Pacific shortfinned eel OMP2
Japanese eel OMP2
Giant mottled eel OMP2
Channel catfish OMP2
Striped catfish OMP2
Bottlenose catfish OMP2
Milkfish OMP2
Denticle herring OMP2
Zebrafish OMP2
Goldfish OMP2
Electric eel OMP2
Red-bellied piranha OMP2
Southern platyfish OMP2
Tongue sole OMP2
Torafugu OMP2
Three-spined stickleback OMP2
Burtons mouthbrooder OMP2
Nile tilapia OMP2
Milkfish OMP1
Zebrafish OMP1
Goldfish OMP1
Southern platyfish OMP1
Tongue sole OMP1
Three-spined stickleback OMP1
Torafugu OMP1
Burtons mouthbrooder OMP1
Nile tilapia OMP1
ompa/OMP2
ompb/OMP1
teleost
OMP
0.1
100
100
79
84
100
100
100
100
74
75
100
100
100
100
73
99
87
100
99
100

Figure S1

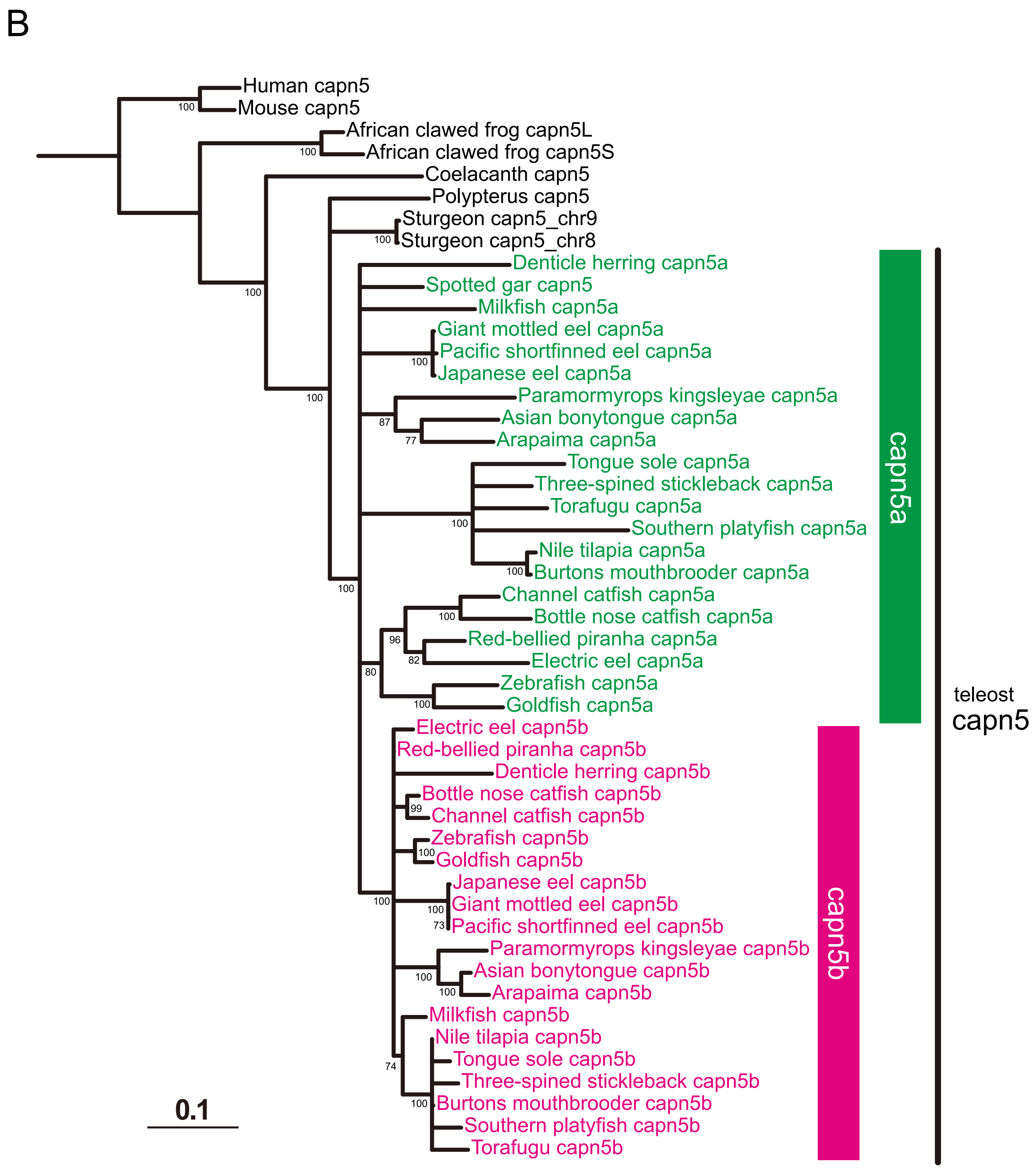
B
Human capn5
Mouse capn5
African clawed frog capn5L
African clawed frog capn5S
Coelacanth capn5
Polypterus capn5
Sturgeon capn5_chr9
Sturgeon capn5_chr8
Denticle herring capn5a
Spotted gar capn5
Milkfish capn5a
Giant mottled eel capn5a
Pacific shortfinned eel capn5a
Japanese eel capn5a
Paramormyrops kingsleyae capn5a
Asian bonytongue capn5a
Arapaima capn5a
Tongue sole capn5a
Three-spined stickleback capn5a
Torafugu capn5a
Southern platyfish capn5a
Nile tilapia capn5a
Burtons mouthbrooder capn5a
Channel catfish capn5a
Bottle nose catfish capn5a
Red-bellied piranha capn5a
Electric eel capn5a
Zebrafish capn5a
Goldfish capn5a
Electric eel capn5b
Red-bellied piranha capn5b
Denticle herring capn5b
Bottle nose catfish capn5b
Channel catfish capn5b
Zebrafish capn5b
Goldfish capn5b
Japanese eel capn5b
Giant mottled eel capn5b
Pacific shortfinned eel capn5b
Paramormyrops kingsleyae capn5b
Asian bonytongue capn5b
Arapaima capn5b
Milkfish capn5b
Nile tilapia capn5b
Tongue sole capn5b
Three-spined stickleback capn5b
Burtons mouthbrooder capn5b
Southern platyfish capn5b
Torafugu capn5b
capn5a
capn5b
teleost
capn5
0.1

Figure S1

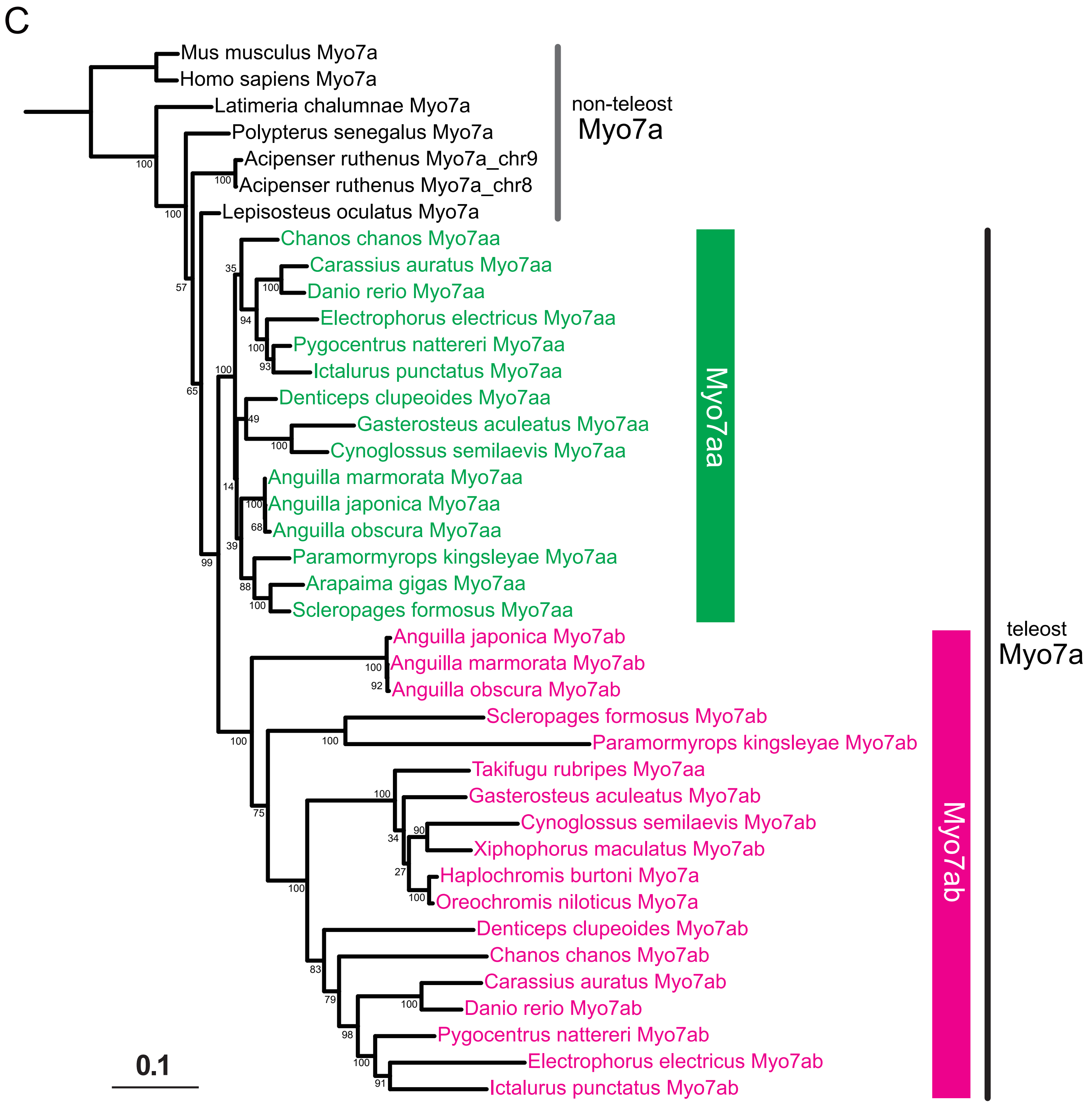
C
Mus musculus Myo7a
Homo sapiens Myo7a
Latimeria chalumnae Myo7a
Polypterus senegalus Myo7a
Acipenser ruthenus Myo7a_chr9
Acipenser ruthenus Myo7a_chr8
Lepisosteus oculatus Myo7a
non-teleost Myo7a
Chanos chanos Myo7aa
Carassius auratus Myo7aa
Danio rerio Myo7aa
Electrophorus electricus Myo7aa
Pygocentrus nattereri Myo7aa
Ictalurus punctatus Myo7aa
Denticeps clupeoides Myo7aa
Gasterosteus aculeatus Myo7aa
Cynoglossus semilaevis Myo7aa
Anguilla marmorata Myo7aa
Anguilla japonica Myo7aa
Anguilla obscura Myo7aa
Paramormyrops kingsleyae Myo7aa
Arapaima gigas Myo7aa
Scleropages formosus Myo7aa
Myo7aa
Anguilla japonica Myo7ab
Anguilla marmorata Myo7ab
Anguilla obscura Myo7ab
Scleropages formosus Myo7ab
Paramormyrops kingsleyae Myo7ab
Takifugu rubripes Myo7aa
Gasterosteus aculeatus Myo7ab
Cynoglossus semilaevis Myo7ab
Xiphophorus maculatus Myo7ab
Haplochromis burtoni Myo7a
Oreochromis niloticus Myo7a
Denticeps clupeoides Myo7ab
Chanos chanos Myo7ab
Carassius auratus Myo7ab
Danio rerio Myo7ab
Pygocentrus nattereri Myo7ab
Electrophorus electricus Myo7ab
Ictalurus punctatus Myo7ab
Myo7ab
teleost Myo7a
0.1

Figure S1

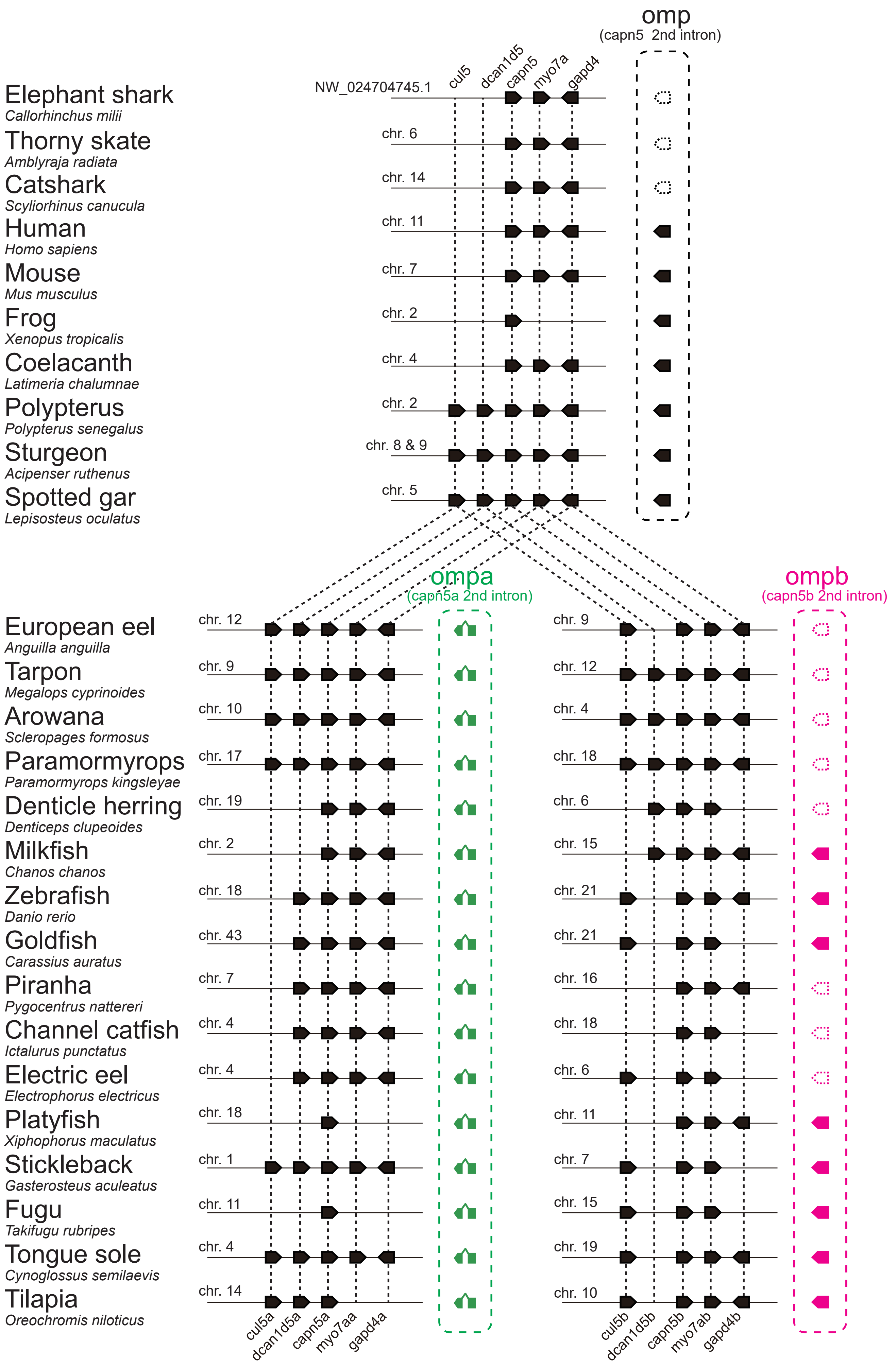
omp
(capn5 2nd intron)
NW_024704745.1
cul5
dcan1d5
capn5
myo7a
gapd4
Elephant shark
Callorhinchus milii
Thorny skate
Amblyraja radiata
chr. 6
Catshark
Scyliorhinus canucula
chr. 14
Human
Homo sapiens
chr. 11
Mouse
Mus musculus
chr. 7
Frog
Xenopus tropicalis
chr. 2
Coelacanth
Latimeria chalumnae
chr. 4
Polypterus
Polypterus senegalus
chr. 2
Sturgeon
Acipenser ruthenus
chr. 8 & 9
Spotted gar
Lepisosteus oculatus
chr. 5
ompa
(capn5a 2nd intron)
ompb
(capn5b 2nd intron)
European eel
Anguilla anguilla
chr. 12
chr. 9
Tarpon
Megalops cyprinoides
chr. 9
chr. 12
Arowana
Scleropages formosus
chr. 10
chr. 4
Paramormyrops
Paramormyrops kingsleyae
chr. 17
chr. 18
Denticle herring
Denticeps clupeoides
chr. 19
chr. 6
Milkfish
Chanos chanos
chr. 2
chr. 15
Zebrafish
Danio rerio
chr. 18
chr. 21
Goldfish
Carassius auratus
chr. 43
chr. 21
Piranha
Pygocentrus nattereri
chr. 7
chr. 16
Channel catfish
Ictalurus punctatus
chr. 4
chr. 18
Electric eel
Electrophorus electricus
chr. 4
chr. 6
Platyfish
Xiphophorus maculatus
chr. 18
chr. 11
Stickleback
Gasterosteus aculeatus
chr. 1
chr. 7
Fugu
Takifugu rubripes
chr. 11
chr. 15
Tongue sole
Cynoglossus semilaevis
chr. 4
chr. 19
Tilapia
Oreochromis niloticus
chr. 14
chr. 10
cul5a
dcan1d5a
capn5a
myo7aa
gapd4a
cul5b
dcan1d5b
capn5b
myo7ab
gapd4b

Figure S2

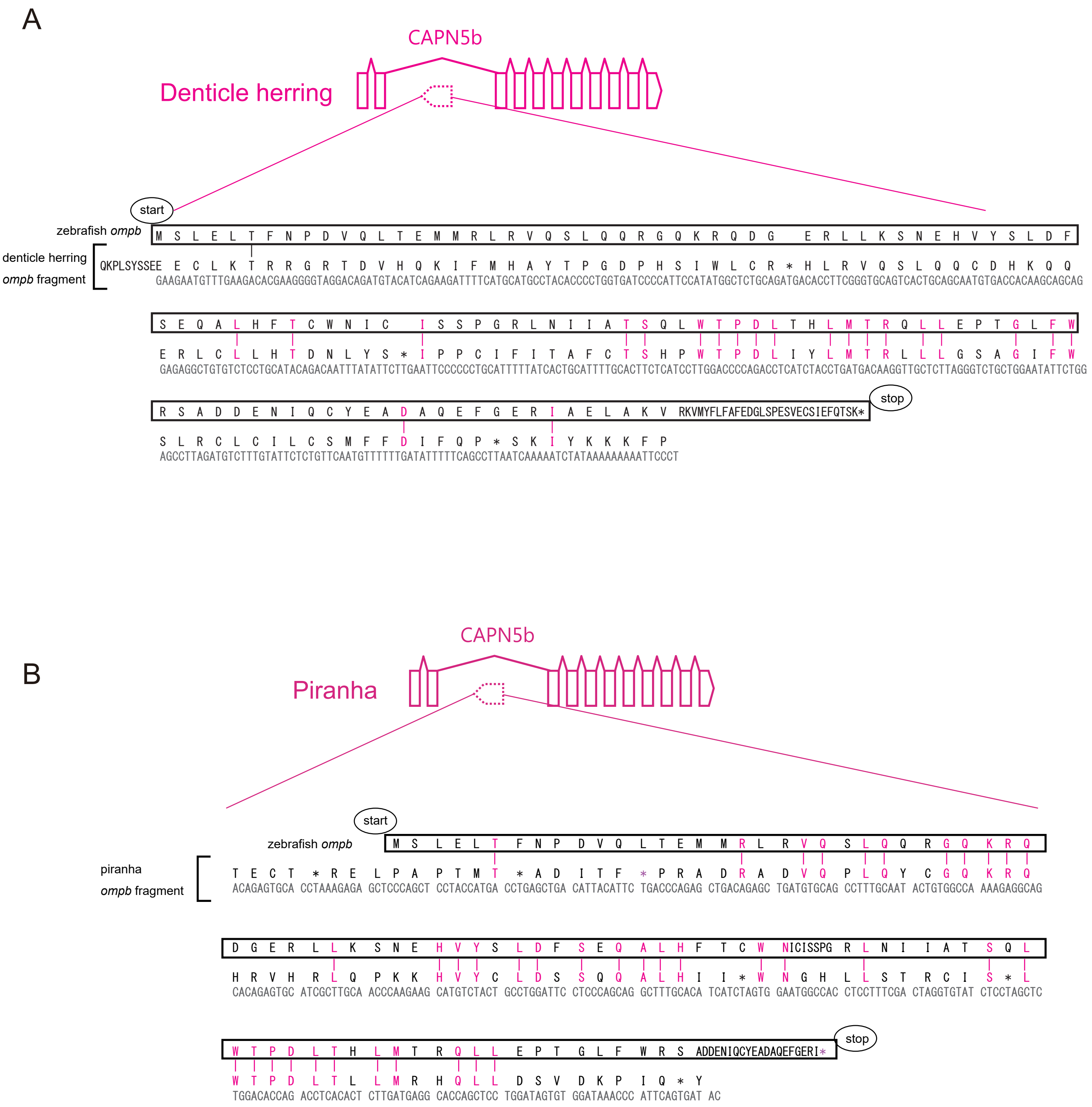
A
CAPN5b
Denticle herring
start
zebrafish ompb
denticle herring
ompb fragment
stop
B
CAPN5b
Piranha
start
zebrafish ompb
piranha
ompb fragment
stop

### Figure S3

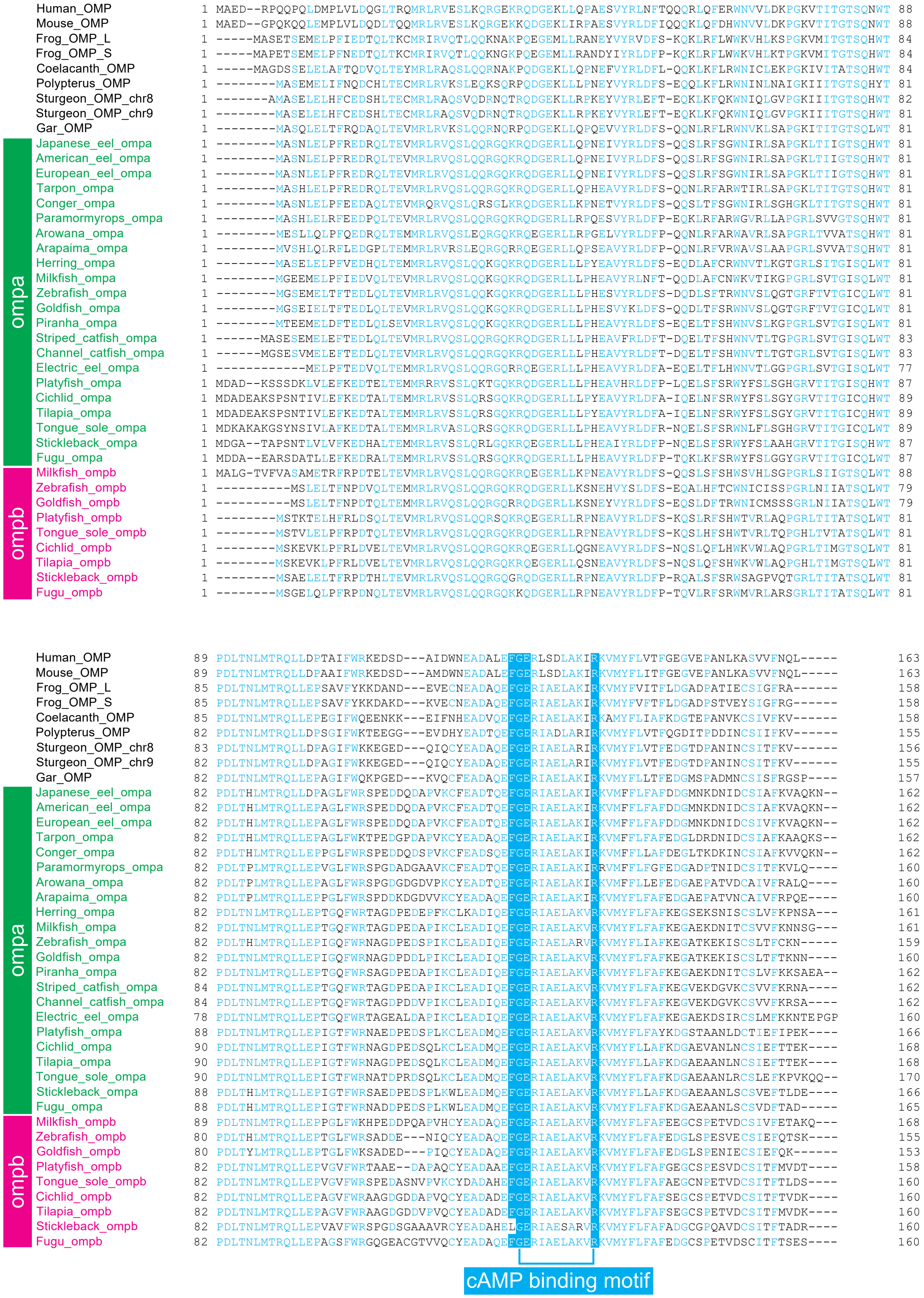
Human_OMP
Mouse_OMP
Frog_OMP_L
Frog_OMP_S
Coelacanth_OMP
Polypterus_OMP
Sturgeon_OMP_chr8
Sturgeon_OMP_chr9
Gar_OMP
Japanese_eel_ompa
American_eel_ompa
European_eel_ompa
Tarpon_ompa
Conger_ompa
Paramormyrops_ompa
Arowana_ompa
Arapaima_ompa
Herring_ompa
Milkfish_ompa
Zebrafish_ompa
Goldfish_ompa
Piranha_ompa
Striped_catfish_ompa
Channel_catfish_ompa
Electric_eel_ompa
Platyfish_ompa
Cichlid_ompa
Tilapia_ompa
Tongue_sole_ompa
Stickleback_ompa
Fugu_ompa
Milkfish_ompb
Zebrafish_ompb
Goldfish_ompb
Platyfish_ompb
Tongue_sole_ompb
Cichlid_ompb
Tilapia_ompb
Stickleback_ompb
Fugu_ompb
ompa
ompb
cAMP binding motif

### Figure S4

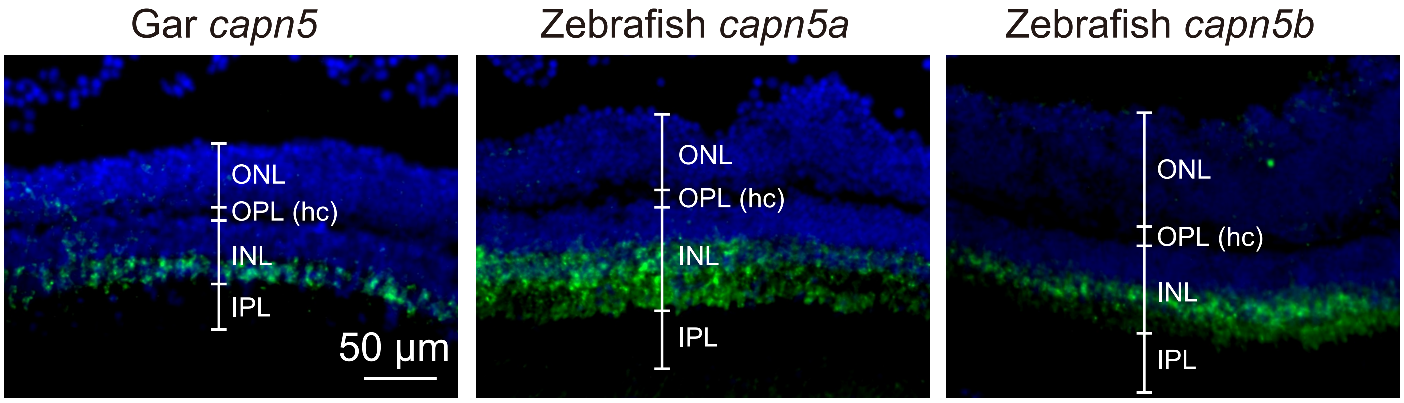

Gar *capn5*
Zebrafish *capn5a*
Zebrafish *capn5b*
ONL
OPL (hc)
INL
IPL
50 µm
ONL
OPL (hc)
INL
IPL
ONL
OPL (hc)
INL
IPL


### Figure S5

### Figure S6

A

B

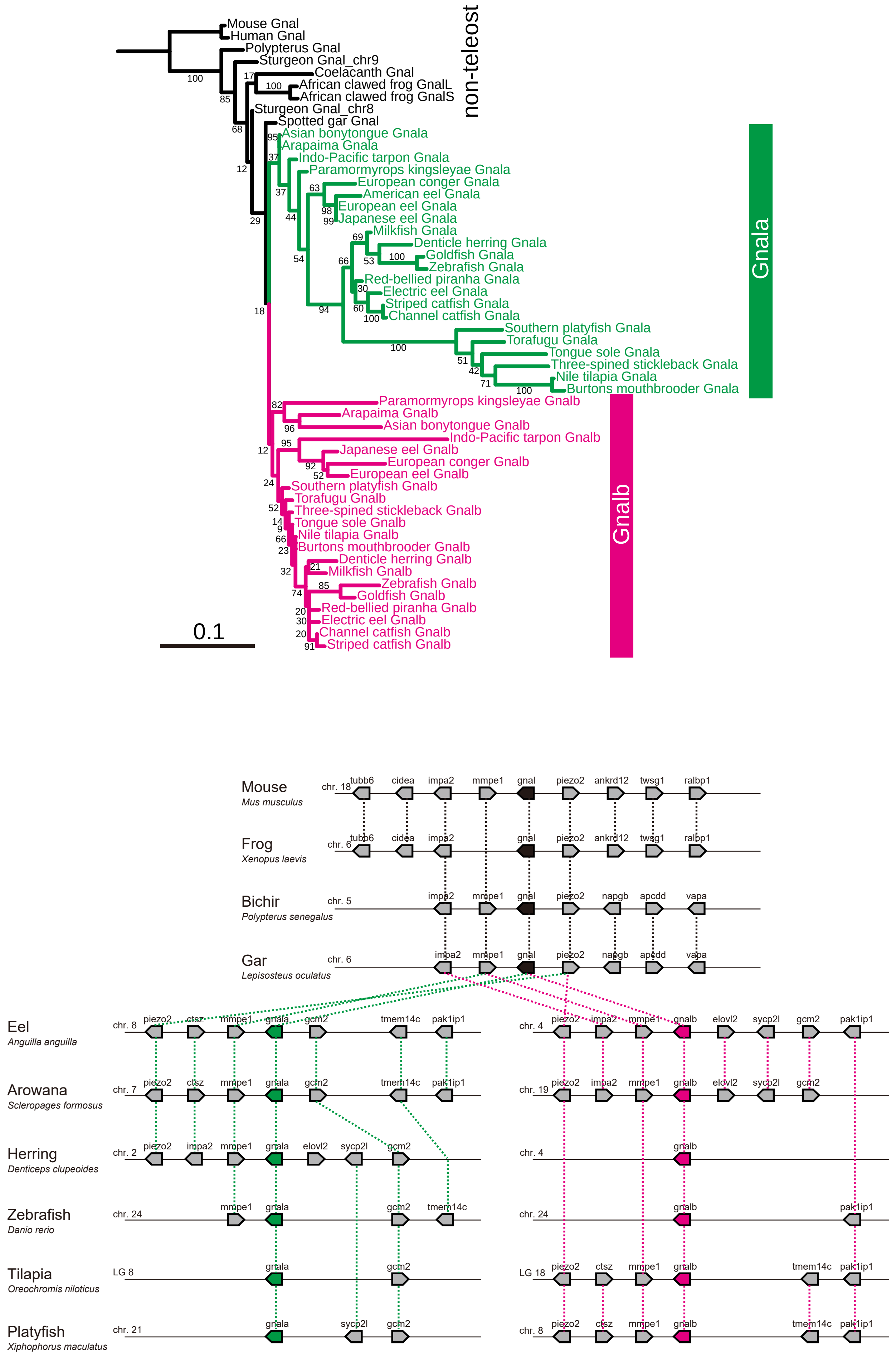
non-teleost
Mouse Gnal
Human Gnal
Polypterus Gnal
Sturgeon Gnal_chr9
Coelacanth Gnal
African clawed frog GnalL
African clawed frog GnalS
Sturgeon Gnal_chr8
Spotted gar Gnal
Asian bonytongue Gnala
Arapaima Gnala
Indo-Pacific tarpon Gnala
Paramormyrops kingsleyae Gnala
European conger Gnala
American eel Gnala
European eel Gnala
Japanese eel Gnala
Milkfish Gnala
Denticle herring Gnala
Goldfish Gnala
Zebrafish Gnala
Red-bellied piranha Gnala
Electric eel Gnala
Striped catfish Gnala
Channel catfish Gnala
Southern platyfish Gnala
Torafugu Gnala
Tongue sole Gnala
Three-spined stickleback Gnala
Nile tilapia Gnala
Burtons mouthbrooder Gnala
Gnala
Paramormyrops kingsleyae Gnalb
Arapaima Gnalb
Asian bonytongue Gnalb
Indo-Pacific tarpon Gnalb
Japanese eel Gnalb
European conger Gnalb
European eel Gnalb
Southern platyfish Gnalb
Torafugu Gnalb
Three-spined stickleback Gnalb
Tongue sole Gnalb
Nile tilapia Gnalb
Burtons mouthbrooder Gnalb
Denticle herring Gnalb
Milkfish Gnalb
Zebrafish Gnalb
Goldfish Gnalb
Red-bellied piranha Gnalb
Electric eel Gnalb
Channel catfish Gnalb
Striped catfish Gnalb
Gnalb
0.1
Mouse
Mus musculus
chr. 18
tubb6 cidea impa2 mmpe1 gnal piezo2 ankrd12 twsg1 ralbp1
Frog
Xenopus laevis
chr. 6
tubb6 cidea impa2 gnal piezo2 ankrd12 twsg1 ralbp1
Bichir
Polypterus senegalus
chr. 5
impa2 mmpe1 gnal piezo2 napgb apcdd vapa
Gar
Lepisosteus oculatus
chr. 6
impa2 mmpe1 gnal piezo2 napgb apcdd vapa
Eel
Anguilla anguilla
chr. 8
piezo2 ctsz mmpe1 gnala gcm2 tmem14c pak1ip1
chr. 4
piezo2 impa2 mmpe1 gnalb elovl2 sycp2l gcm2 pak1ip1
Arowana
Scleropages formosus
chr. 7
piezo2 ctsz mmpe1 gnala gcm2 tmem14c pak1ip1
chr. 19
piezo2 impa2 mmpe1 gnalb elovl2 sycp2l gcm2
Herring
Denticeps clupeoides
chr. 2
piezo2 impa2 mmpe1 gnala elovl2 sycp2l gcm2
chr. 4
gnalb
Zebrafish
Danio rerio
chr. 24
mmpe1 gnala gcm2 tmem14c
chr. 24
gnalb pak1ip1
Tilapia
Oreochromis niloticus
LG 8
gnala gcm2
LG 18
piezo2 ctsz mmpe1 gnalb tmem14c pak1ip1
Platyfish
Xiphophorus maculatus
chr. 21
gnala sycp2l gcm2
chr. 8
piezo2 ctsz mmpe1 gnalb tmem14c pak1ip1

### Figure S6

C

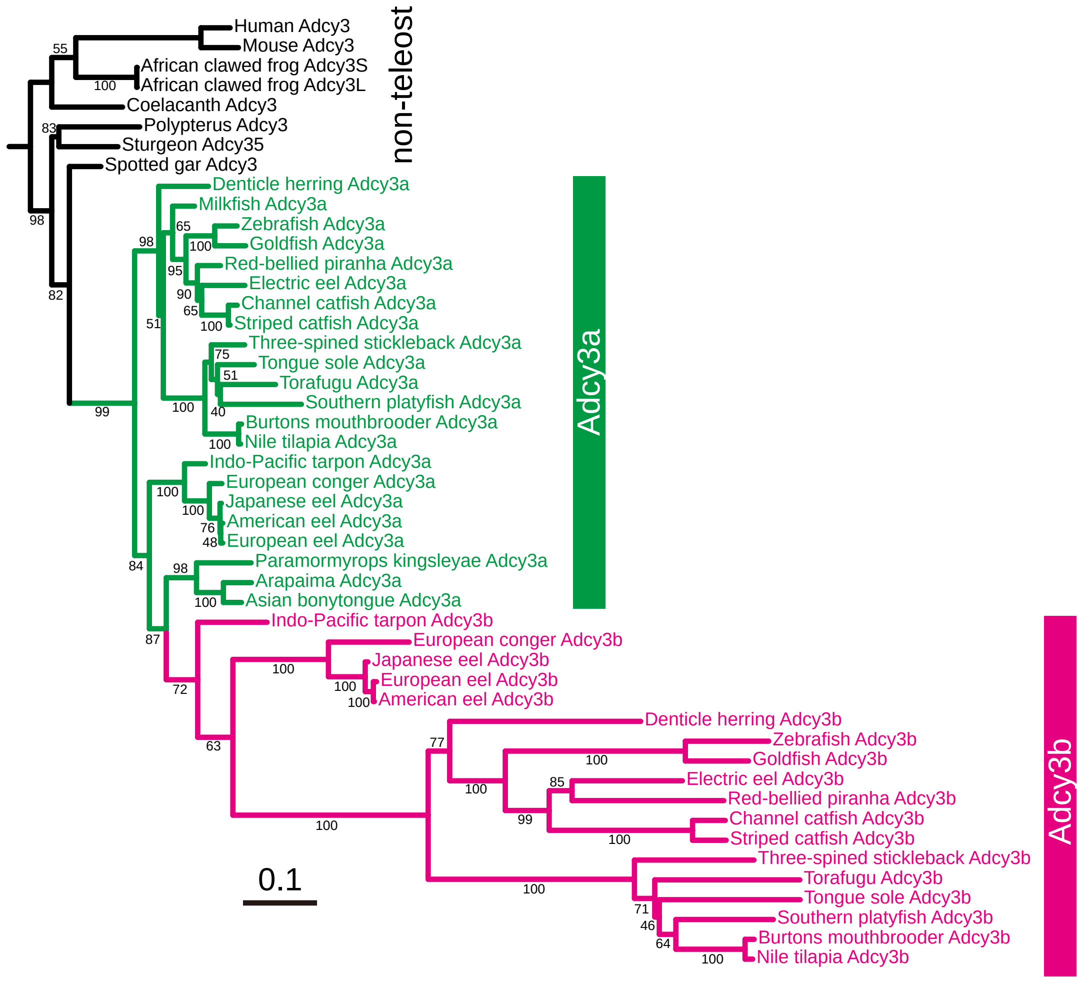
Human Adcy3
Mouse Adcy3
African clawed frog Adcy3S
African clawed frog Adcy3L
Coelacanth Adcy3
Polypterus Adcy3
Sturgeon Adcy35
Spotted gar Adcy3
non-teleost
Denticle herring Adcy3a
Milkfish Adcy3a
Zebrafish Adcy3a
Goldfish Adcy3a
Red-bellied piranha Adcy3a
Electric eel Adcy3a
Channel catfish Adcy3a
Striped catfish Adcy3a
Three-spined stickleback Adcy3a
Tongue sole Adcy3a
Torafugu Adcy3a
Southern platyfish Adcy3a
Burtons mouthbrooder Adcy3a
Nile tilapia Adcy3a
Indo-Pacific tarpon Adcy3a
European conger Adcy3a
Japanese eel Adcy3a
American eel Adcy3a
European eel Adcy3a
Paramormyrops kingsleyae Adcy3a
Arapaima Adcy3a
Asian bonytongue Adcy3a
Adcy3a
Indo-Pacific tarpon Adcy3b
European conger Adcy3b
Japanese eel Adcy3b
European eel Adcy3b
American eel Adcy3b
Denticle herring Adcy3b
Zebrafish Adcy3b
Goldfish Adcy3b
Electric eel Adcy3b
Red-bellied piranha Adcy3b
Channel catfish Adcy3b
Striped catfish Adcy3b
Three-spined stickleback Adcy3b
Torafugu Adcy3b
Tongue sole Adcy3b
Southern platyfish Adcy3b
Burtons mouthbrooder Adcy3b
Nile tilapia Adcy3b
Adcy3b
0.1

D

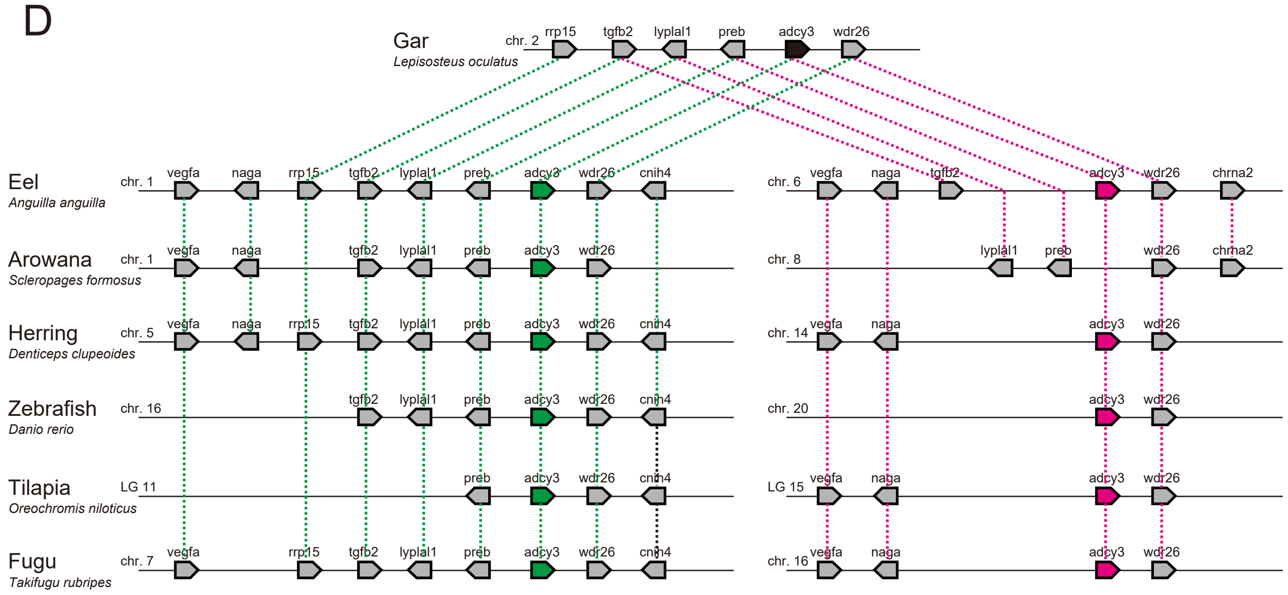
Gar
*Lepisosteus oculatus*
chr. 2
rrp15
tgfb2
lyplal1
preb
adcy3
wdr26
Eel
*Anguilla anguilla*
chr. 1
chr. 6
Arowana
*Scleropages formosus*
chr. 8
Herring
*Denticeps clupeoides*
chr. 5
chr. 14
Zebrafish
*Danio rerio*
chr. 16
chr. 20
Tilapia
*Oreochromis niloticus*
LG 11
LG 15
Fugu
*Takifugu rubripes*
chr. 7
chr. 16
vegfa
naga
cnih4
chrna2

### Figure S6

E

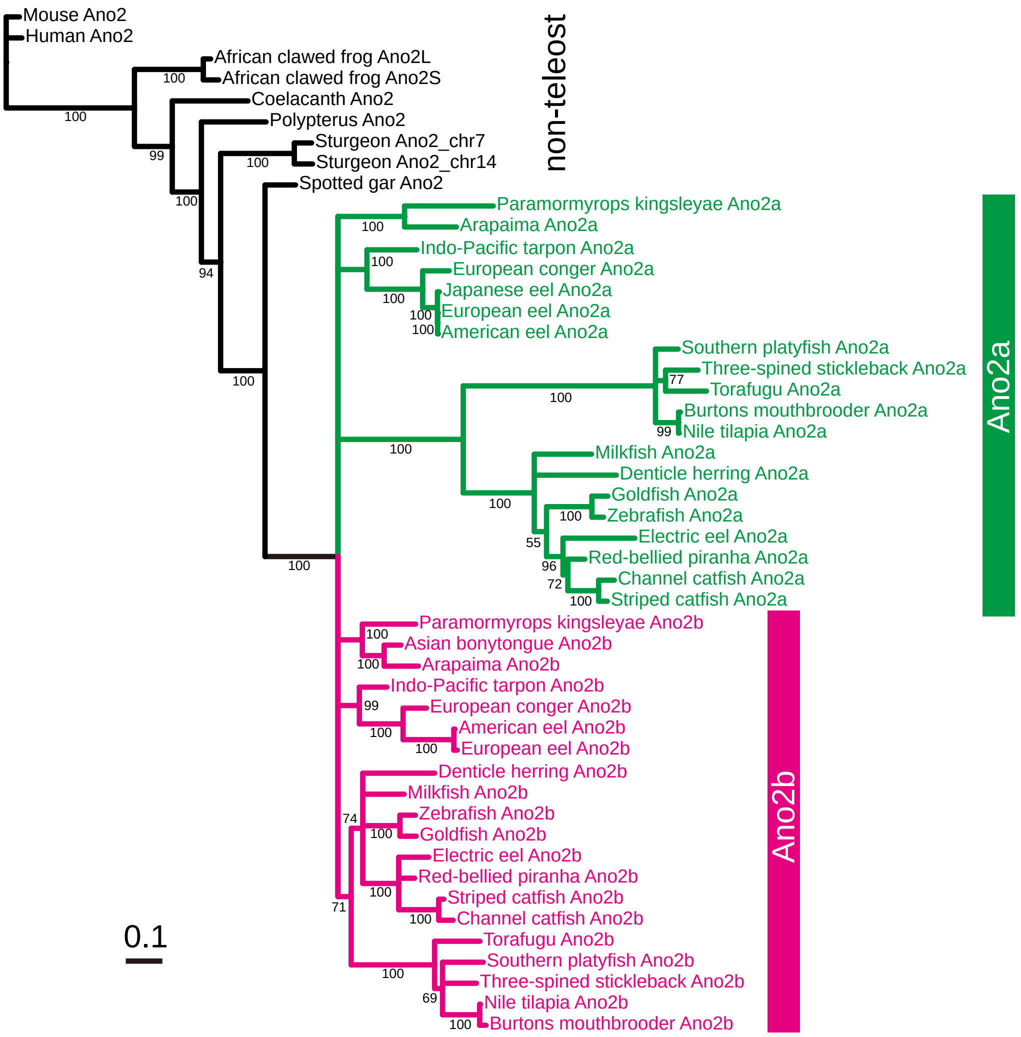
Mouse Ano2
Human Ano2
African clawed frog Ano2L
African clawed frog Ano2S
Coelacanth Ano2
Polypterus Ano2
Sturgeon Ano2_chr7
Sturgeon Ano2_chr14
Spotted gar Ano2
non-teleost
Paramormyrops kingsleyae Ano2a
Arapaima Ano2a
Indo-Pacific tarpon Ano2a
European conger Ano2a
Japanese eel Ano2a
European eel Ano2a
American eel Ano2a
Southern platyfish Ano2a
Three-spined stickleback Ano2a
Torafugu Ano2a
Burtons mouthbrooder Ano2a
Nile tilapia Ano2a
Milkfish Ano2a
Denticle herring Ano2a
Goldfish Ano2a
Zebrafish Ano2a
Electric eel Ano2a
Red-bellied piranha Ano2a
Channel catfish Ano2a
Striped catfish Ano2a
Ano2a
Paramormyrops kingsleyae Ano2b
Asian bonytongue Ano2b
Arapaima Ano2b
Indo-Pacific tarpon Ano2b
European conger Ano2b
American eel Ano2b
European eel Ano2b
Denticle herring Ano2b
Milkfish Ano2b
Zebrafish Ano2b
Goldfish Ano2b
Electric eel Ano2b
Red-bellied piranha Ano2b
Striped catfish Ano2b
Channel catfish Ano2b
Torafugu Ano2b
Southern platyfish Ano2b
Three-spined stickleback Ano2b
Nile tilapia Ano2b
Burtons mouthbrooder Ano2b
Ano2b
0.1

F

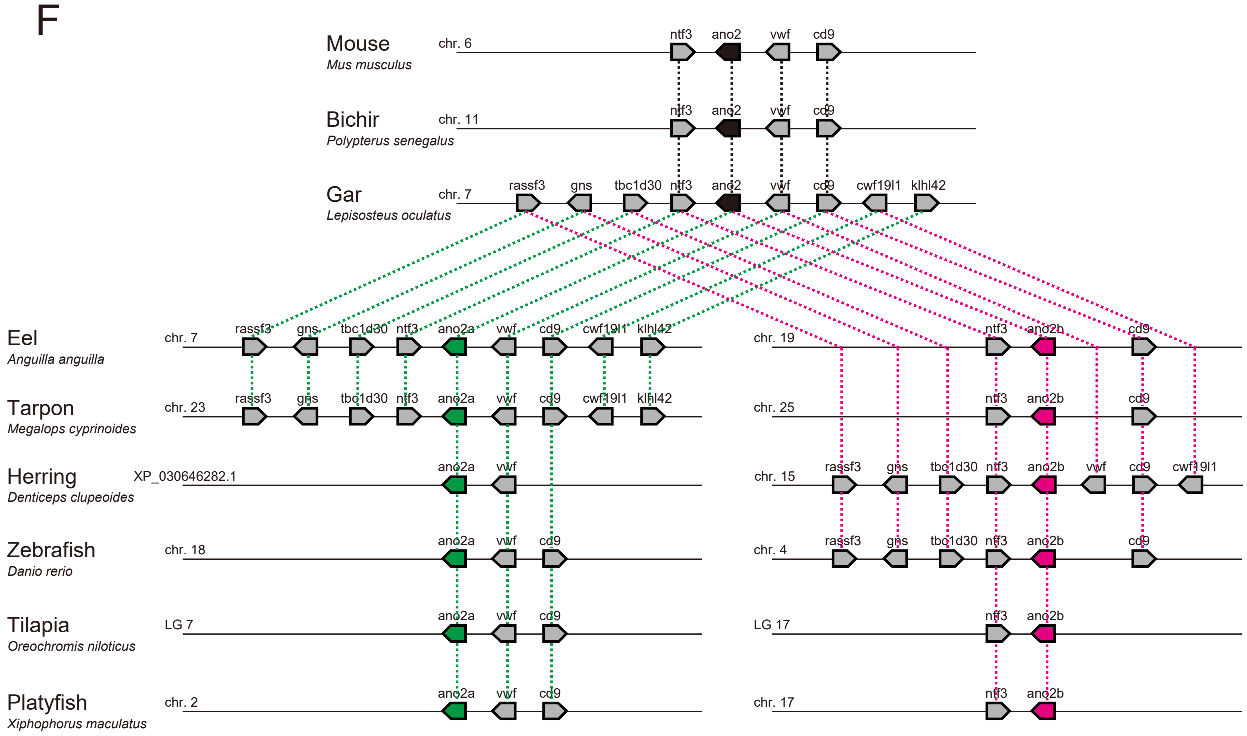
Mouse
Mus musculus
chr. 6
Bichir
Polypterus senegalus
chr. 11
Gar
Lepisosteus oculatus
chr. 7
rassf3 gns tbc1d30 ntf3 ano2 vwf cd9 cwf19l1 klhl42
Eel
Anguilla anguilla
chr. 7
chr. 19
Tarpon
Megalops cyprinoides
chr. 23
chr. 25
Herring
Denticeps clupeoides
XP_030646282.1
chr. 15
Zebrafish
Danio rerio
chr. 18
chr. 4
Tilapia
Oreochromis niloticus
LG 7
LG 17
Platyfish
Xiphophorus maculatus
chr. 2
chr. 17
ano2a
ano2b

### Figure S6

G

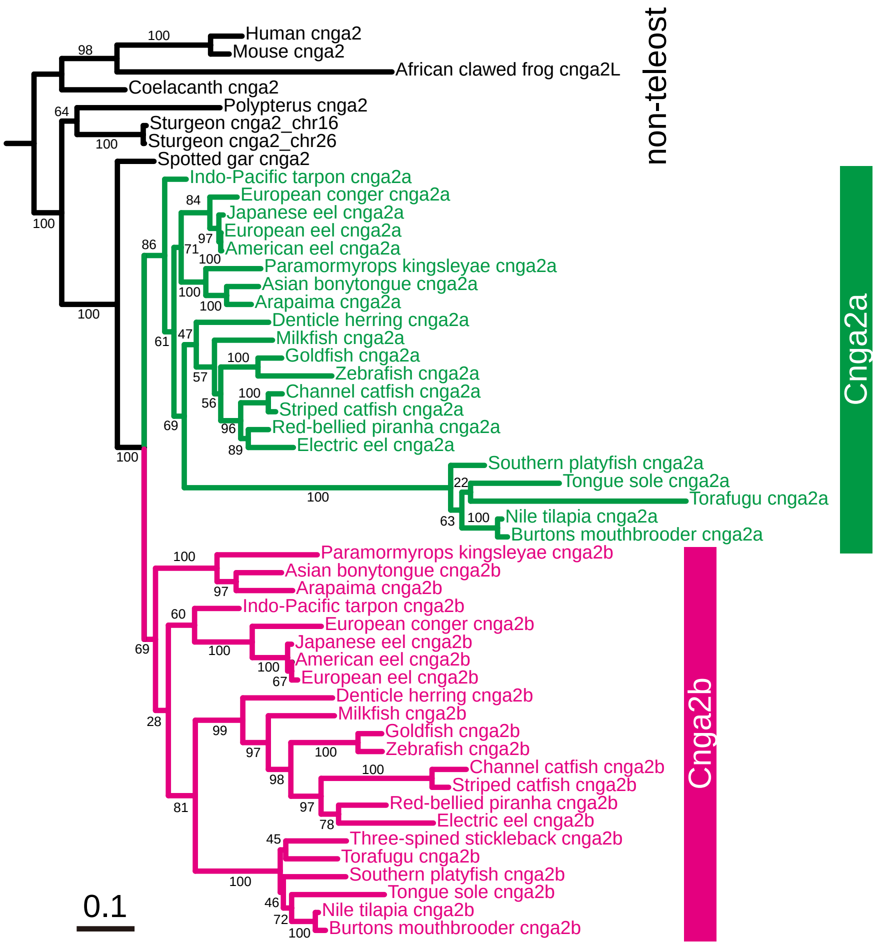

Human cnga2
Mouse cnga2
African clawed frog cnga2L
Coelacanth cnga2
Polypterus cnga2
Sturgeon cnga2_chr16
Sturgeon cnga2_chr26
Spotted gar cnga2
Indo-Pacific tarpon cnga2a
European conger cnga2a
Japanese eel cnga2a
European eel cnga2a
American eel cnga2a
Paramormyrops kingsleyae cnga2a
Asian bonytongue cnga2a
Arapaima cnga2a
Denticle herring cnga2a
Milkfish cnga2a
Goldfish cnga2a
Zebrafish cnga2a
Channel catfish cnga2a
Striped catfish cnga2a
Red-bellied piranha cnga2a
Electric eel cnga2a
Southern platyfish cnga2a
Tongue sole cnga2a
Torafugu cnga2a
Nile tilapia cnga2a
Burtons mouthbrooder cnga2a
Paramormyrops kingsleyae cnga2b
Asian bonytongue cnga2b
Arapaima cnga2b
Indo-Pacific tarpon cnga2b
European conger cnga2b
Japanese eel cnga2b
American eel cnga2b
European eel cnga2b
Denticle herring cnga2b
Milkfish cnga2b
Goldfish cnga2b
Zebrafish cnga2b
Channel catfish cnga2b
Striped catfish cnga2b
Red-bellied piranha cnga2b
Electric eel cnga2b
Three-spined stickleback cnga2b
Torafugu cnga2b
Southern platyfish cnga2b
Tongue sole cnga2b
Nile tilapia cnga2b
Burtons mouthbrooder cnga2b
non-teleost
Cnga2a
Cnga2b
0.1


H

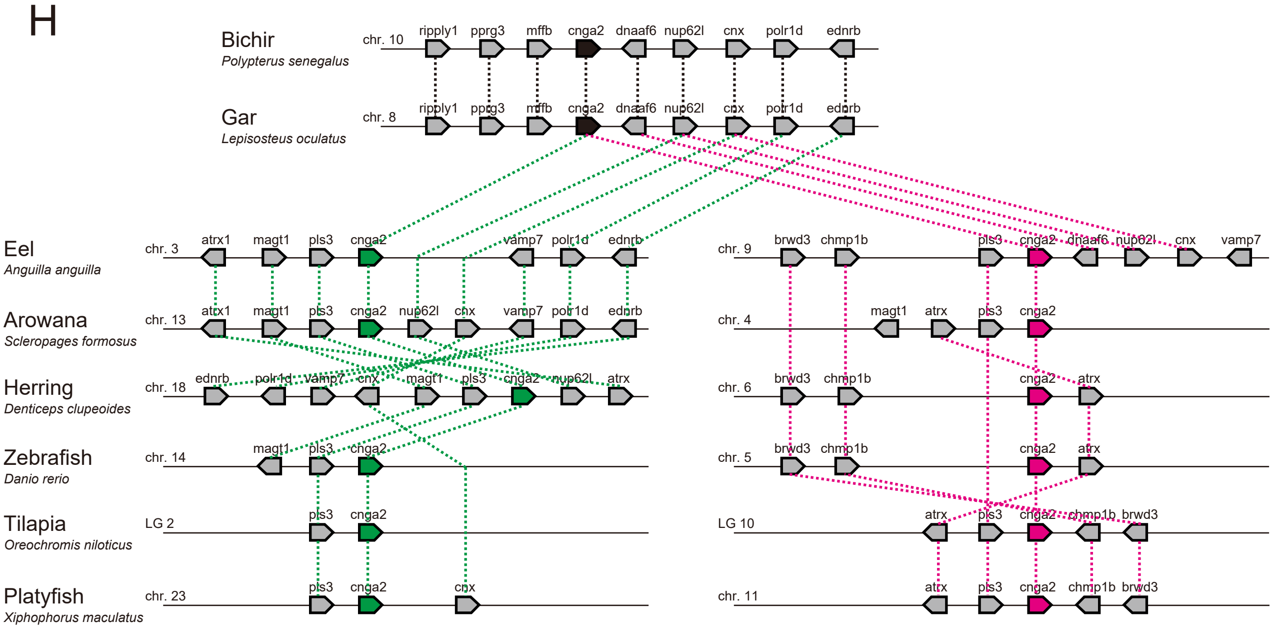

Bichir
Polypterus senegalus
chr. 10
Gar
Lepisosteus oculatus
chr. 8
ripply1 pprg3 mffb cnga2 dnaaf6 nup62l cnx polr1d ednrb
Eel
Anguilla anguilla
chr. 3
chr. 9
Arowana
Scleropages formosus
chr. 13
chr. 4
Herring
Denticeps clupeoides
chr. 18
chr. 6
Zebrafish
Danio rerio
chr. 14
chr. 5
Tilapia
Oreochromis niloticus
LG 2
LG 10
Platyfish
Xiphophorus maculatus
chr. 23
chr. 11
atrx1 magt1 pls3 cnga2 vamp7 polr1d ednrb
brwd3 chmp1b pls3 cnga2 dnaaf6 nup62l cnx vamp7
atrx1 magt1 pls3 cnga2 nup62l cnx vamp7 polr1d ednrb
magt1 atrx pls3 cnga2
ednrb polr1d vamp7 cnx magt1 pls3 cnga2 nup62l atrx
brwd3 chmp1b cnga2 atrx
magt1 pls3 cnga2
brwd3 chmp1b cnga2 atrx
pls3 cnga2
atrx pls3 cnga2 chmp1b brwd3
pls3 cnga2 cnx
atrx pls3 cnga2 chmp1b brwd3

### Figure S6

I

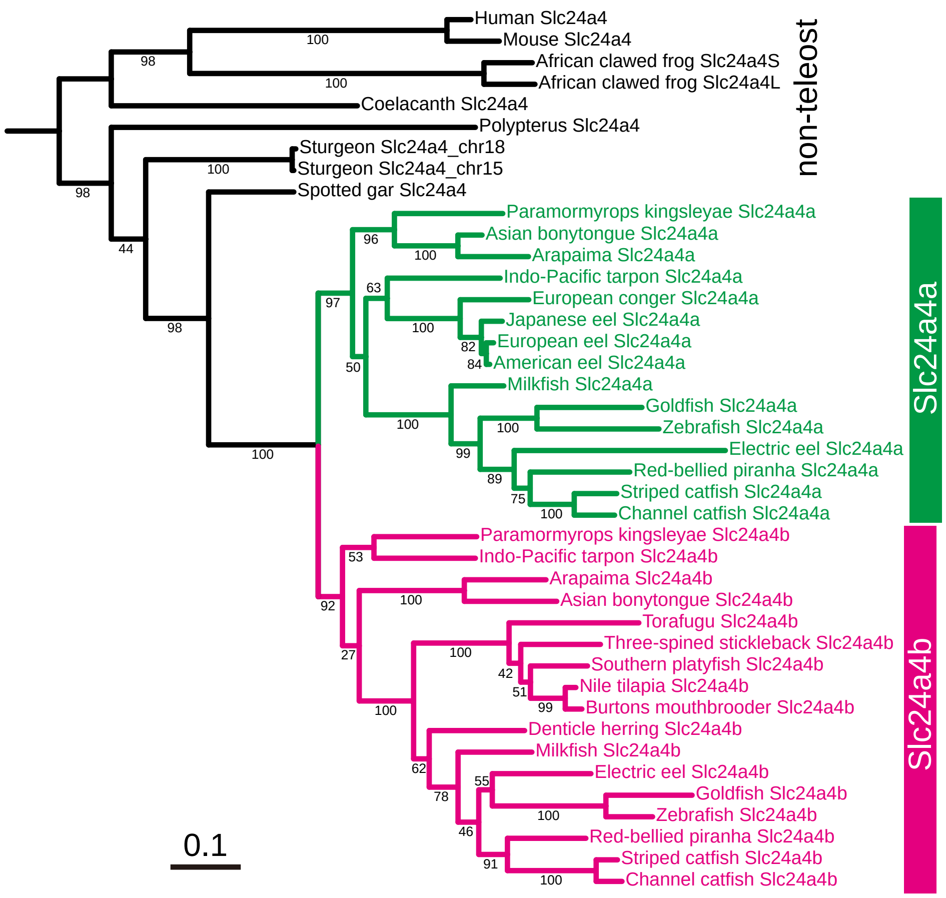

Human Slc24a4
Mouse Slc24a4
African clawed frog Slc24a4S
African clawed frog Slc24a4L
Coelacanth Slc24a4
Polypterus Slc24a4
Sturgeon Slc24a4_chr18
Sturgeon Slc24a4_chr15
Spotted gar Slc24a4
Paramormyrops kingsleyae Slc24a4a
Asian bonytongue Slc24a4a
Arapaima Slc24a4a
Indo-Pacific tarpon Slc24a4a
European conger Slc24a4a
Japanese eel Slc24a4a
European eel Slc24a4a
American eel Slc24a4a
Milkfish Slc24a4a
Goldfish Slc24a4a
Zebrafish Slc24a4a
Electric eel Slc24a4a
Red-bellied piranha Slc24a4a
Striped catfish Slc24a4a
Channel catfish Slc24a4a
Paramormyrops kingsleyae Slc24a4b
Indo-Pacific tarpon Slc24a4b
Arapaima Slc24a4b
Asian bonytongue Slc24a4b
Torafugu Slc24a4b
Three-spined stickleback Slc24a4b
Southern platyfish Slc24a4b
Nile tilapia Slc24a4b
Burtons mouthbrooder Slc24a4b
Denticle herring Slc24a4b
Milkfish Slc24a4b
Electric eel Slc24a4b
Goldfish Slc24a4b
Zebrafish Slc24a4b
Red-bellied piranha Slc24a4b
Striped catfish Slc24a4b
Channel catfish Slc24a4b
non-teleost
Slc24a4a
Slc24a4b
0.1


J

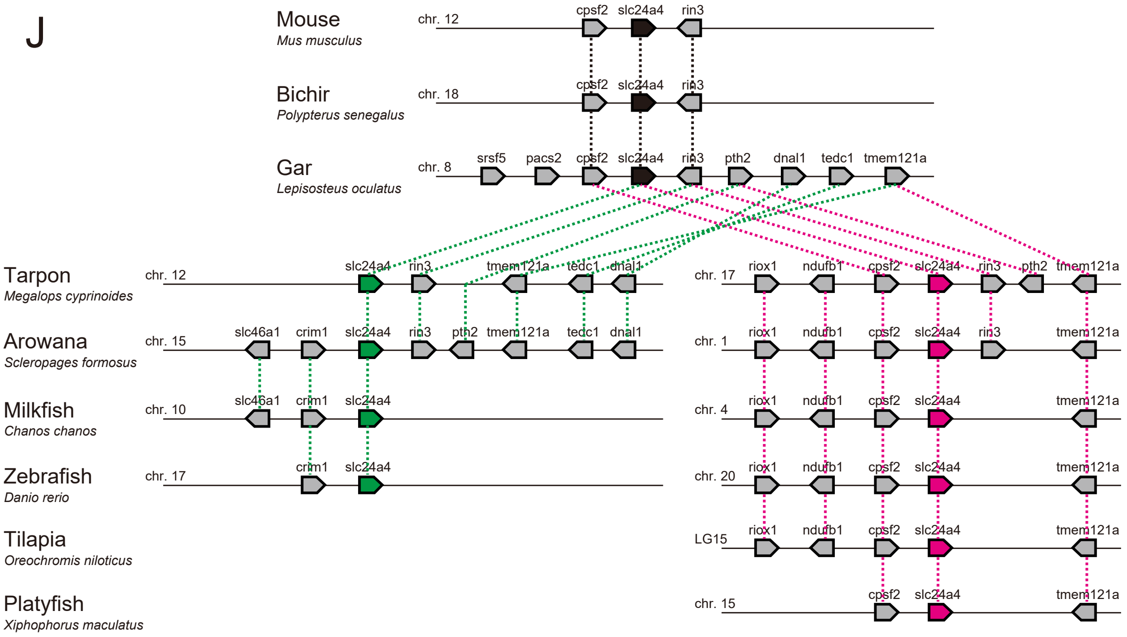

Mouse
Mus musculus
Bichir
Polypterus senegalus
Gar
Lepisosteus oculatus
Tarpon
Megalops cyprinoides
Arowana
Scleropages formosus
Milkfish
Chanos chanos
Zebrafish
Danio rerio
Tilapia
Oreochromis niloticus
Platyfish
Xiphophorus maculatus
chr. 12
chr. 18
chr. 8
chr. 15
chr. 10
chr. 17
chr. 1
chr. 4
chr. 20
LG15
chr. 15
cpsf2
slc24a4
rin3
srsf5
pacs2
pth2
dnal1
tedc1
tmem121a
slc46a1
crim1
riox1
ndufb1

>DreOmpa_upstream
GAGGTCGACGGTATCGATAAGCTTTCTAGAAAGTGTTGAAGAAATGGTTAACA
GACCAAACGTTTTGCAGTGGCCTAGCCAGAGTCCTGACTTTAACCCAACTGAG
AATCTGTAGAGGAAGCTAATGCAATTCATCTTTATTTTTATAGTGCTTTTATAA
GATTATGTAGATTATGGCAAAGCAGCTTCACATAGAAGATTATAGTGATAAAGA
TCAGGCTGATGGTAAACTCTACAACCTAAAAGAGTAGAAGCTCATCGATAAAG
ATAAATGGGCAAAAATACAAGTGGAGACATGCAGAAAGCAGCTCAGCAATTAT
AGGATGTGTTTGATTGCTATAAATAGCCAAAACAGGATATTGATCATTGAAAA
GGGTATGAATAATTTTGGACAGGACACTTTTTGTTCAAATGTAAATAAAATCT
GTGAAATTAGTTGGTTTTTTCCACAACGATGACTCTTGTCAACGTCTTATTAT
CTTTTGGGAGAAGCCAGTGTCATATTTGGTCAAAAACAACAACTTGCATCAGA
ATTATTCAACACCAGAATTTGCTTAACTGTATATATTCCAGCTAAACAAAAAGT
AGTAAAACCTTCTCCAGAAGACAAATACAACAGGACATTTTGTTATATATTAGG
GGTGTCAAAATTAATTGTCGGTGCACCGCGATGCAGACGCGGACAATTCGGT
ATTGATTCAGTAATAACCATAACCGGTTATTATGTAGTGACGTCATTTATCTCC
TATGCGCTCTGTCGCGAGGGAGGTGAGCGCCATTATTTACAACACTCAGCCA
ACTTGGGGCCGGCCCGTGGCATAGGCACCATAGGCAAATGCTTAGGGCGCTG
TATATCCAGTGGGGCGCCAGAAATGAGCGCGCTTCAGTTGGTTTTGTTTTTG
ATTTTCCTGACACACATTCAGTTATAGACGGCAATAACTCAAAAAAACGCTTAC
CCTAAAAAGATCTAAAGTGTTTCCTACAAAAAGACAAAGCTGGTTATGTTTCA
CAGCTGTCTTCGCTCGACATTACGAGCACTGCTCCGTTTAAGACCTCTCAGTC
TGCGTTTACTTTAGCCGGCAGAGCTCGCGCTGAGAGGAGCTCTCTGTAAGGG
ACCGTCGGAAAAGTGCTTATTTTTTTCGCTGCTCCGCTAAGCGCAAGCTCTC
CCTGTTGCAAGGGCTTAAGTGTGTTTGTTTTTGTTTGTTTGATGCTGTCTAT
GTTTGTGAATGTGTTTGAATGAGAGACAGTGTGTTGCTGTGTGTCTGTGTGT
TTATACAGACAGCATGTTATAGCCTCCCCCCTAAAATATAACACTATATATGGA
AAGCATCGTCAATGCACCGTGATGCACCAAGATATGGAATTGAACCGAATCGA
TGGCATGATAATCGTAACCGAACCGAACTGTGAGACAAGTATAGGTTCACACC
TCGGAAATTGTGAATAAAATTACAATTTTACCTCAAATGCACAAAACAAAGCAA
AACAGAAAGTCTGGAAGTTTTCCAACAGCAGAAAGCATCTGTTGCCTAGAGCT
GCAACACTGAATCTGTTTGCTTTGCTCCTCCTCCTTTTTCCAAAACTAAAGTC
GAGTCTCAGAGAATCAATAAAGTCCCCATCTGAGCGAGTGTGTGTGCCCGAG
GGGACGCAGTGGCGCTGCAGCGGATGGAAAAGTGAAGGCTGAGCGAGGAAA
AGCTTCATTCACACACAGAGAGAATCAGCAAAACAGCATCCCGCGGGATCCA
CCGGTAAGTCGCCACCA

### Figure S7
